# Cerebellar transcranial AC stimulation produces frequency and phase-dependent changes in Purkinje cell complex spike waveform

**DOI:** 10.64898/2026.11.12.766205

**Authors:** Devry Mourra, Jack Sloane, Mesut Sahin, Eric J. Lang

## Abstract

Cerebellar transcranial alternating current stimulation (ctACS) can improve motor learning and may speed rehabilitation. The mechanisms by which ctACS induces long-term changes in brain function that underlie these effects are not known, and knowledge of them could greatly improve the application of ctACS. One potential mechanism is alteration of Purkinje cell (PC) complex spike (CS) activity, as CS activity is a key regulator of cerebellar synaptic plasticity. Thus, we investigated how ctACS modifies CS firing rates and waveform. Extracellular recordings were obtained from cerebellar PCs in anesthetized adult rats while ctACS (250 μA) was applied across a range of frequencies (0.5 - 50 Hz). The effect of ctACS on CS waveform was quantified via measuring changes in the overall size of the CS and in the number and amplitude of the spikelets that comprise a CS. ctACS produced phase-dependent modulation of the CS waveform in a majority of PCs at most tested frequencies, with 1 Hz stimulation causing modulation in all PCs tested at the frequency. The modulation reflected changes within the normal range of CS waveforms rather than an expansion of that range. Analysis of individual spikelet periods (windows) revealed that ctACS modified discrete portions of the CS, with 93.8% of PCs showing significant modulation in at least one window. Notably, the strength of phase-dependent CS waveform modulation was relatively consistent across frequencies, in contrast to the frequency dependence of simple spike (SS) entrainment. Significant modulation of spikelet amplitude was found more often than that of spikelet number; however, the relative strength of the modulation when present was similar for the two parameters. CS waveform modulation was correlated with SS entrainment strength and was strongest in PCs located near the cerebellar surface (i.e., near the apices of the folia).

Together, these findings demonstrate that ctACS exerts phase-dependent control over CS waveform. Moreover, the differing frequency dependence of the waveform effects and SS firing modulation suggests that it may be possible to target CS-related plasticity selectively, providing a potential strategy for selectively modulating cerebellar learning with ctACS.

---

Electrical neuromodulation techniques are increasingly being used to treat a range of neurological and psychiatric disorders (Grimaldi et al., 2014; Priori et al., 2014; Fertonani et al., 2015; Grimaldi et al., 2016). In particular, several lines of evidence suggest that cerebellar transcranial AC stimulation (ctACS) can not only improve motor learning but also affect cognitive and emotional functions in both normal and brain-injured individuals (Naro et al., 2017; Miyaguchi et al., 2018; Ahn et al., 2019; Feng et al., 2024; Kaiser et al., 2025). These beneficial effects imply that ctACS affects processes, such as synaptic plasticity, that underlie long-term changes in brain function. Currently, we have little to no knowledge of which of these processes ctACS might affect, and such knowledge would be valuable for guiding our use of ctACS. To begin addressing this gap, we investigated how ctACS affects Purkinje cell (PC) complex spike (CS) activity, as CSs have long been implicated in the control of synaptic plasticity in the cerebellum (Ito, 2001).

ctACS could affect CS activity in at least two ways that would impact plasticity. First, it could modulate CS firing rate. CSs are triggered by activation of the PC by its climbing fiber (terminal portion of the olivocerebellar axon). Given that the inferior olive (IO), the origin of olivocerebellar pathway, is in the ventral medulla, ctACS is unlikely to modulate the activity of IO neurons via a direct polarization of IO neurons, because of the IO’s distance from the ctACS electrode. However, ctACS strongly modulates PC simple spike (SS) activity (Asan and Sahin, 2019a; Mourra et al., 2025), and ctACS induced changes in SS activity could affect the IO via the disynaptic PC to cerebellar nuclei to IO pathway. Consistent with this possibility, pharmacological and optogenetic manipulations of PC SS activity have been shown to alter CS activity in the same cerebellar cortical region (Marshall and Lang, 2009; Chaumont et al., 2013).

A second possible mechanism by which ctACS could influence synaptic plasticity is altering the waveform of the CS. There is evidence that the type and strength of plasticity induced by CSs vary with CS waveform (i.e., changes in spikelet number) (Mathy et al., 2009; Rasmussen et al., 2013; Yang and Lisberger, 2014), but see (Titley et al., 2020). In addition, changes in CS waveform have also been correlated with motor learning (Yang and Lisberger, 2014). Thus, understanding the effect of ctACS on CS waveform could provide clues as to whether ctACS can influence CS-mediated plasticity.

Here, we report that ctACS induces phase-modulated changes in CS waveform across a wide frequency range, reflecting alterations in both the number and amplitude of spikelets comprising a CS. Furthermore, the results indicate that the effects of ctACS on CS waveform have a distinct frequency dependence from its effects on SS activity, suggesting a way to preferentially control different aspects of cerebellar activity.

## Materials and Methods

Experiments were performed in accordance with the National Institute of Health *Guide for the Care and Use of Laboratory Animals*. Experimental protocols were approved by the Institutional Animal Care and Use Committee of New York University School of Medicine.

### Surgical procedures

Non-survival surgeries were performed on adult female Sprague-Dawley rats (273 ± 25.8 g, n = 22). Anesthesia was induced by an injection of ketamine (100 mg/kg) and xylazine (8 mg/kg) intraperitoneally. After the animal reached a surgical plane of anesthesia, a tracheotomy was performed to enable mechanical ventilation, if needed, and a femoral vein catheter was inserted to deliver supplemental anesthesia, which was given as a continuous intravenous injection of a ketamine-xylazine solution (ketamine, 6 μg/kg/min; xylazine, 1 μg/kg/min). The depth of anesthesia was maintained such that pedal reflexes were absent. Temperature was measured using a rectal probe and maintained at 37 °C with the aid of a heating pad (TR-200, Fine Science Tools, Foster City, CA). Animals were placed in a stereotaxic frame, an incision in the skin overlying the dorsal skull was made, and soft tissue was cleared from the dorsal skull surface. Next, a burr hole (∼1mm diameter, centered at AP: -11 mm, LAT: 3.8 mm, relative to Bregma) was made to allow access for a microelectrode to record PC activity in the lateral hemisphere.

Neuronal activity was amplified with either a Dagan amplifier (Model EX1, 500-1000x gain, low and high cutoff filters of 300 Hz and 10 kHz) or an A-M systems (Sequim, WA USA) AC amplifier (Model 1700 with 1000x gain and 300 Hz and 10 kHz cutoff filters), and was recorded using a multichannel recording system (MultiChannel Systems MCS GmbH, Reutlingen, Germany) with a sampling rate of 25 kHz/channel.

### Recording and stimulation procedures

ctACS was applied using Ag/AgCl electrodes, with one placed on the skull surface overlying crus1 on the same side as the recording electrode and posteromedial to the burr hole (relative to bregma, AP: -13.2 mm, LAT: 2.4 mm; Fig. 1a). The other stimulation electrode was inserted below the skin of the opposite shoulder. Each electrode was made from a chlorided silver wire (diameter 0.38 mm, #7830, A-M Systems, Sequim, WA, USA). The end of the electrode placed on the skull was shaped into a 2 mm diameter coil disk and soaked in electrolytic gel for better contact (Spectra 360 Electrode Gel #12-02).

**Fig. 1.**
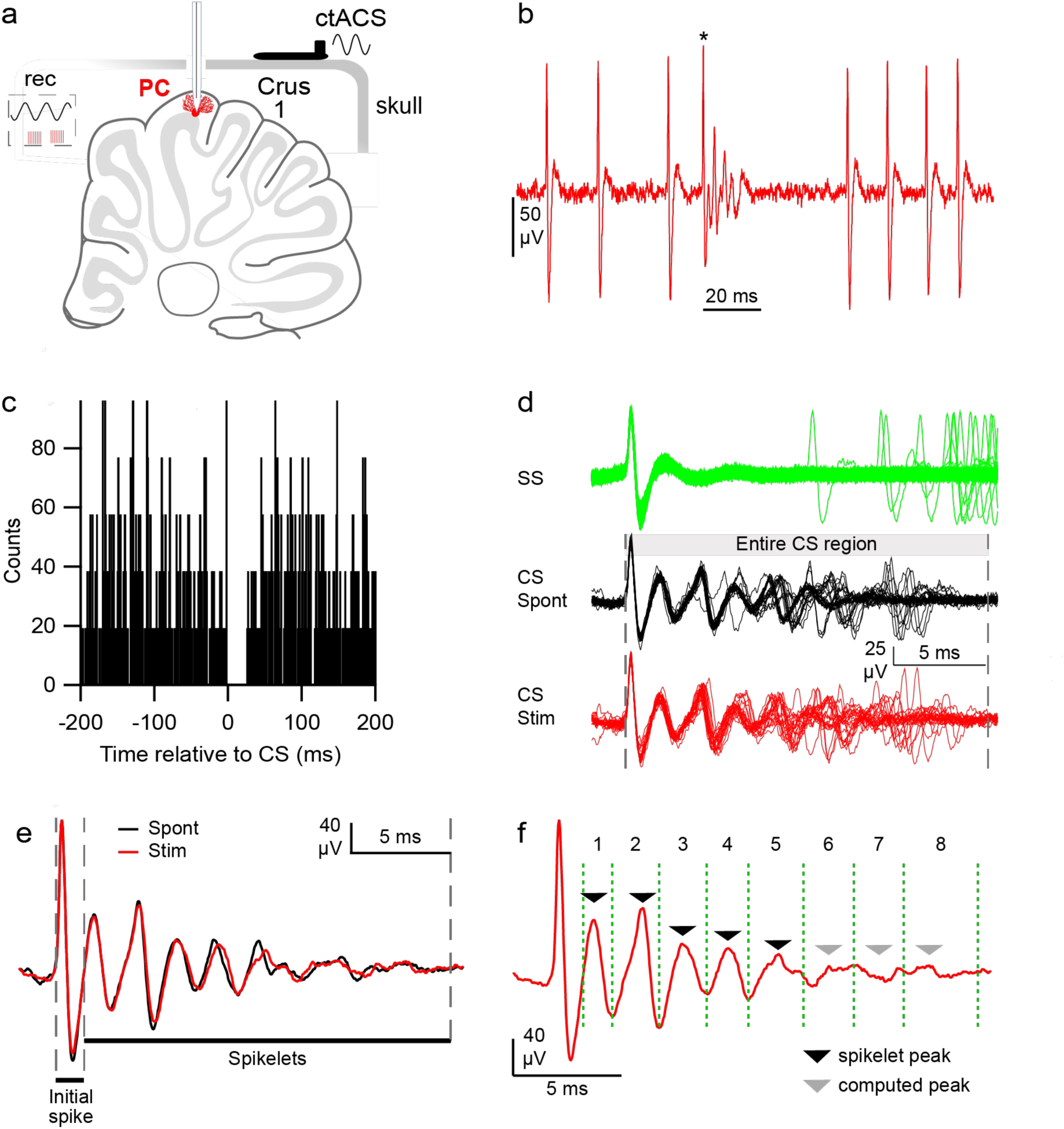
Identification of SS activity and division of CS waveform for analysis. **(a)** Sagittal schematic of the cerebellum and rat skull showing placement of ctACS electrode directly on the skull above the lateral crus I. Recording electrode inserted through a burr hole into the cerebellar cortex. **(b)** Extracellular recording of PC showing a CS (denoted by an asterisk) and SSs. Note the post-CS pause in SSs. **(c)** The peri-CS histogram displays a clear pause in SS firing after a CS (same PC as 1b). **(d)** Overlaid waveforms of SSs (green) and CS waveforms during spontaneous (black) and ctACS (red). The shaded rectangle indicates the entire CS period used for analyses. **(e)** Mean CS waveforms with main analysis divisions indicated: IS period (between first and second dashed lines) and SpkLts period (between second and third dashed lines). **(f)** Mean CS waveform during ctACS (red trace, same as in **e**) showing subdivision of SpkLts period. Windows were defined to surround each detectable peak (black triangles) and computed peaks (gray triangles) in the average waveform. Vertical, dashed, green lines indicate window limits. For details of peak detection and window limit settings, see Methods.

Following the placement of the stimulation electrodes, a glass microelectrode filled with 1 M NaCl was systematically lowered through the burr hole into the cerebellar mass to obtain extracellular recordings of PC activity (DV: 0−3 mm) using a joystick-controlled micromanipulator (Burleigh Inchworm LSS-1000). The recording ground was placed under the skin on the ipsilateral shoulder.

The AC stimulus peak-to-peak amplitude was 500 μA for all recordings. The AC frequency was systematically increased in steps from 0.5 Hz to 50 Hz. When a single unit was isolated, spiking activity was recorded for a 4-minute period for each stimulus frequency, during which AC stimulation at the particular frequency was interleaved with periods of recording with no stimulation (10 seconds on/10 seconds off), allowing for a comparison of activity during the two conditions. AC stimuli were generated by an isolated current stimulator (Model SYS-A395D, WPI, Sarasota, FL) driven by a waveform generator (Model 185, Wavetek, San Diego, CA, USA). The following nominal frequencies spanning the delta to gamma frequency range were tested: 0.5, 1, 5, 10, 20, and 50 Hz. Since the waveform generator had an analog frequency selector knob, stimulation frequency varied slightly from the nominal frequencies across recordings. Overall, the percent changes of actual stimulation frequencies for each nominal frequency were: 0.5 Hz = ± 0.45 %, 1 Hz = ± 1.02 %, 5 Hz = ± 0.61 %, 10 Hz = ± 1.05 %, 20 Hz = ± 0.41 %, and 50 Hz = ± 0.56 %. For simplicity, the nominal frequencies are listed in the text and figure axis labels; however, for statistical analyses in which stimulation frequency was an independent variable, the exact stimulation frequency was used.

### Data Analysis

Data files containing the extracellular recording traces, AC stimulus wave, and trigger times marking the start of the stimulation periods were converted to binary or text files using MC_Datatool software (MultiChannel Systems MCS GmbH), and then imported into Igor Pro 9 (Wavemetrics, Lake Oswego, OR, USA; RRID:SCR_000325). Data analysis was performed using built-in and custom routines with Igor Pro, Microsoft Excel, and R Project for Statistical Computing (RRID:SCR_001905).

To eliminate artifacts due to the AC stimulus, the raw recordings were high-pass filtered using a finite impulse response (FIR) filter with a Hanning window and a cutoff frequency of 250-300 Hz (number of coefficients: 701). Potential spikes were then detected using two voltage windows (upper and lower), with a time limit of 1 ms for crossing within both windows. Principal component analysis (PCA) was applied to the detected waveforms, and a cluster plot of the first two components was used to separate SS and CS waveforms. The set of CSs was then manually reviewed to confirm that it contained only CSs. Waveform analyses were performed in Igor Pro 9 using custom-written routines. PC SSs were identified as such by showing a pause following a CS (Fig. 1b, c). To analyze the CS waveform, a segment of the extracellular recording was extracted from 10 ms before to 30 ms after the onset of each CS (Fig. 1d).

### Defining CS components and calculation of the absolute area under the curve (AAUC)

The AAUC was used to quantify how ctACS affects the CS waveform and its components. It was calculated by rectifying the CS waveform and then summing the absolute value of the area between the curve and zero across all deflections. For analysis purposes, we initially divided the CS into two parts: the initial spike (IS) and spikelet (SpkLt) parts (Fig. 1e), and the AAUC was calculated for each one. The IS period started at the initial deflection of the CS, continued through the first negative and positive peaks (or vice versa), and concluded at the subsequent baseline crossing point (0 mV). The SpkLt period started at the end of the IS period. The endtime of the SpkLt period was first approximated by visual inspection of the average CS waveforms during the spontaneous and stimulated activity periods. Next, the individual traces of all CSs from a PC were overlaid, and the end of the SpkLt period was set such that all SpkLts from all of the CSs concluded prior to that time. The SpkLt period duration varied between 10 and 30 ms across the PC population. To exclude any contamination by SSs, only traces with clear post-CS SS pauses were analyzed.

For some analyses, the SpkLt region was further divided into subregions referred to as ‘SpkLt windows’ to estimate the AAUCs of individual SpkLts. This division was based on the average CS waveform of a PC: peaks within the SpkLt region were detected based on a preset voltage threshold level for each PC. The troughs between each pair of detected peaks were used to set the limits of most of the SpkLt windows, with the following exceptions. For the first SpkLt window, the start of the SpkLt period was used as its beginning limit. For the last detected SpkLt peak, the ending limit of its window was set by adding one half the interspikelet interval between it and the prior SpkLt peak except when that time exceeded the endtime of the entire CS, in which case the CS endtime was used. Usually the window of the last detected SpkLt in the average waveform ended well before the endtime of the CS (i.e., more than the interspikelet interval between the last two detected SpkLt peaks), because SpkLts toward the end of the CS were less aligned and therefore tended to cancel out in the average. When this occurred, the peak times were defined by the interspikelet interval between the two prior peaks: multiples of this interval were added until the completion of the CS. The limits of these windows were then set as the times of these peaks ± one half of the interspikelet interval or the endtime of the CS, whichever came first (Fig. 1f, gray arrowheads).

### Computing phasic modulation in AAUC

To assess the phasic modulation of AAUC for the IS and SpkLt regions, and for the SpkLt windows, we fit a sine wave to the corresponding AAUC values as a function of the phase with respect to the AC cycle, *y* = *y*₀ + *A sin*(*θ* − *φ*). The 95% confidence interval (CI) for *A* was used to determine whether there was statistically significant modulation. If the 95% CI for *A* did not include 0, then the modulation was considered significant. Since the size of the CS (and each SpkLt) varied between cells, we divided *A* by y_0_ to obtain the fractional modulation in waveform size (normalized *A*) to allow comparisons across the population. The phase of each value was taken as the time of the beginning limit of its window.

### Calculation of the circular mean vector of PC activity during ctACS

The circular mean vector (CMV) was used as a non-parametric measure of the modulation of firing rate (i.e., degree of entrainment to the ctACS stimulation). To calculate the CMV, every spike was represented as a unit vector in the direction equal to its phase with respect to the AC cycle and summed vectorially. This sum was then divided by the total number of spikes during the stimulation to obtain the CMV. The magnitude of the CMV (r̅) ranges from 0 to 1 and measures the strength of modulation, where 1 indicates that the activity is perfectly phase-locked to the ctACS stimulation, and 0 indicates an entirely uniform distribution of phases (i.e., no phase-locking). The CMV’s phase (t̅) provides a measure of the timing of spike responses relative to the AC stimulation.

### Detection and measurement of spikelets

For each CS waveform, spikelets were identified using a peak-to-trough detection algorithm; local minima and maxima were sequentially detected within the spikelet period, and candidate events were defined by peaks occurring within a specified peak-to-trough time interval. Spikelets were accepted when the amplitude difference between the peak and adjacent troughs exceeded a predefined threshold. Spikelet amplitude was measured as the peak-to-trough difference to minimize false positives arising from baseline fluctuations. The output was then manually reviewed.

Spikelet amplitude was calculated as the maximum amplitude, defined as the greater of the differences between the peak and the two surrounding troughs; values were then averaged to provide the mean spikelet amplitude for each CS.

### Statistics

Unless otherwise stated, t-tests, paired or unpaired, or ANOVA, as appropriate, were used to test for statistical significance based on the built-in functions in R and libraries for circular statistics (Hankin, 2015). Mixed models were chosen to handle unbalanced data and to treat frequency as a continuous variable. We modeled the cell as a random effect across frequency. A linear mixed-effects model (LMM) was used with the lme4 (RRID:SCR_015654) (Bates et al., 2015), to calculate Wald chi-square or *F* test values using lmerTest (Kuznetsova et al., 2017).

To assess whether the CMV magnitude (r̅) was statistically significant, we used the Rayleigh test for uniformity for circular data (r̅ > 0, *p* < 0.05). r̅ values are presented with their circular standard deviation 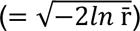 in the text (Mardia, 1975). Except for circular data, means are reported with their standard deviation (SD). Correlations for linear data were calculated using Pearson’s *r*. For phase angle correlations, the Jammalamadaka-Sarma Coefficient, *ρ_JS_*, was used.

### Histological and stereotaxic procedures for identifying the locations of PCs

After a recording track was completed, alcian blue dye was injected in the middle (depth ∼2 mm) and the end of the track (depth ∼4 mm). At the conclusion of the experiment, the animal was perfused under deep anesthesia with normal saline, followed by 10% formalin. The brain was removed, soaked in 10% formalin for at least 1-2 days, and then transferred to a 30% sucrose/10% formalin solution until it sank. Parasagittal 60-μm sections of the cerebellum were prepared on a freezing microtome and mounted on chrome–alum gelatin-coated slides for counterstaining with cresyl violet. To identify PC locations, tissue shrinkage was calculated by measuring the distance between the two dye spots and then taking the ratio of the measured distance to the difference in the micrometer readings for the dye spots. Micrometer values for the depths of the recorded cells were then adjusted by this factor and plotted along the track. Typically, tissue shrinkage was ∼10%.

In experiments with multiple tracks, dye spots were placed only on some tracks to avoid ambiguity in defining each track. In these cases, when cells were recorded on unmarked tracks, their location was defined using shrinkage-corrected micrometer readings and comparison to the nearest dye-labeled track.

## Results

To begin assessing potential mechanisms by which ctACS can induce long-lasting changes in cerebellar function, we investigated whether it could alter CS firing rates and waveform. Recordings were made from 49 cerebellar cortical cells during ctACS in adult anesthetized rats (n = 15), located in either the simplex or crus I lobules, based on their recording depths and histological controls. Of these, 30 were identified as PCs by the presence of CS activity, along with SS activity, the latter indicated by a post-CS pause in activity (Fig. 1b, c). The presence of large amplitude SSs indicated that the recordings were from locations near or at the PC soma. From this PC population, 16 PCs had clearly separable CS and SS activity (i.e., a minimum post-CS pause in SS activity that was longer in duration than the longest CS of that PC) and a large signal-to-noise ratio that allowed the SpkLts of the CSs to be clearly distinguished from baseline noise (Fig. 1b, d). For these 16 PCs, CS activity (firing rate or waveform) and/or SS activity were modified by ctACS at least at one stimulus frequency, with SS modulation providing an independent measure that the PC experienced an electric field capable of significantly modulating its activity.

### Changes in average and phase-dependent CS firing rates

ctACS did not alter the average CS firing rate (spontaneous, 0.28 ± 0.11 Hz; ctACS, 0.71 ± 1.61 Hz; *t*_(14)_ = 1.04, *p =* 0.318). Phase-dependent modulation of CS firing rate was observed in only a minority of PCs at each tested frequency (0.5 Hz: 1/15, 1.0 Hz: 1/10, 5 Hz: 3/8, 10 Hz: 3/8, 20 Hz: 1/9, and 50 Hz: 0/9; Rayleigh test, *p* < 0.05). In addition, only 1 PC was significantly modulated across the entire 0.5 - 20 Hz frequency range. Because of the relative rarity of CS firing rate modulation induced by ctACS at the tested intensity, the remainder of the results will focus on the effects of ctACS on CS waveform. However, it is worth noting that CS firing rate modulation was much more common with direct surface AC stimulation (Avlar et al. unpublished results).

### ctACS has limited effect on the average CS waveform

To evaluate the impact of ctACS on CS waveform, the total duration of the CS waveform of each PC was defined as the longest observed waveform of all of its individual CSs during both the spontaneous activity and ctACS periods (Fig. 1d, middle and lower traces). The waveform was then divided into IS and SpkLt regions using the mean CS waveform (Fig. 1e, dashed lines) as described in the Methods, and the AAUC of each individual CS was then calculated for each of these regions.

We first investigated whether applying ctACS caused a change in the average AAUC values of the CS from those of spontaneous CSs. At the individual PC level, only a minority of PCs demonstrated significant changes in AAUC for the SpkLt region from spontaneous values at any tested stimulus frequency, with the percentage of PCs showing a change ranging from 11.1 - 37.5%: 0.5 Hz: 26.7% (4/15), 1.0 Hz: 20% (2/10), 5 Hz: 12.5% (1/8), 10 Hz: 37.5% (3/8), 20 Hz: 22.2% (2/9), and 50 Hz: 11.1% (1/9). The percentage of PCs that showed a significant steady state change across all tested frequencies was 12.5% (2/16), and the percentage of PCs that had such a change for at least one frequency was 50% (8/16). However, the direction of the change was inconsistent across PCs, and thus at the population level no significant difference was found in the normalized mean AAUC between spontaneous CSs and those occurring during ctACS (Fig. 2a; Table 1). Also, ctACS generally did not affect the limits of the AAUC range, as in most cases there was no significant change in the average minimum AAUC or the average maximum AAUC (Fig. 2b, c; Table 1). Similarly, minimal changes were found for the IS portion (data not shown).

**Fig. 2.**
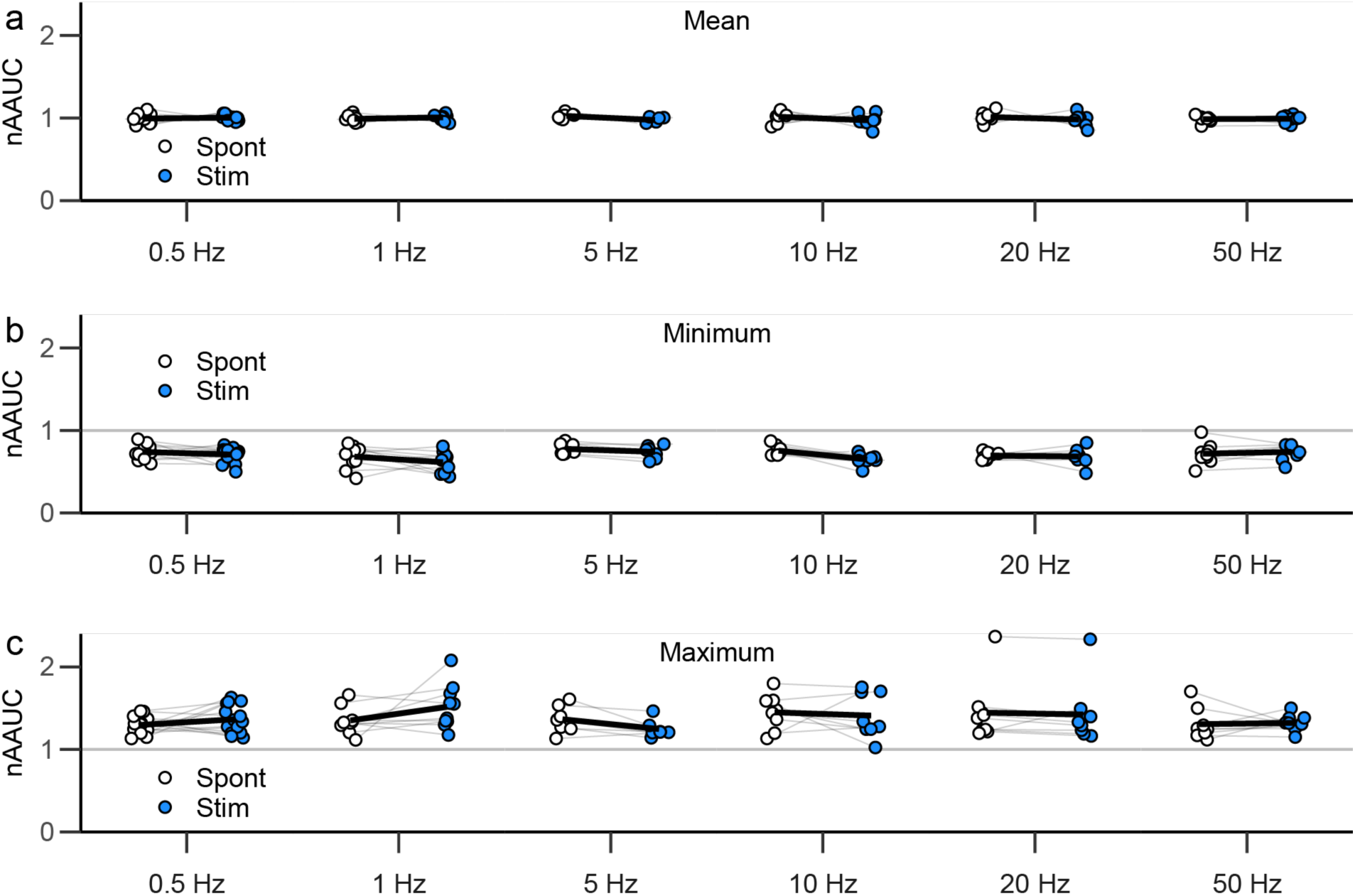
ctACS does not induce steady-state changes in CS waveform. (a-c) Pair plots, comparing average normalized AAUC **(a)**, minimum normalized AAUC values **(b)**, and maximum normalized AAUC values **(c)** during spontaneous activity (Spont) and stimulation (Stim) at each frequency for the SpkLt region. Each circle show the value for an individual PC, each gray line connects the spontaneous and stimulation data for the same PC. Population averages are shown by the black lines.

**Table 1.**
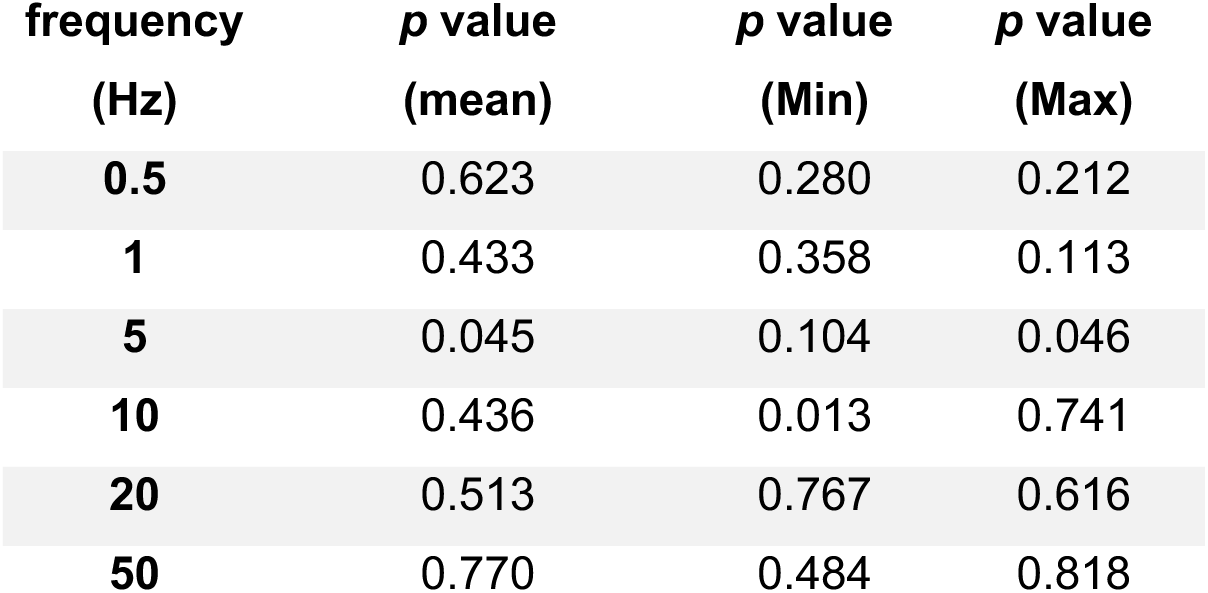
P-values for changes in CS normalized AAUC parameters across the PC population. The listed p-values are uncorrected for multiple tests. If a Hommel correction for testing the PCs at 6 frequencies is applied, none are significant at the 0.05 level.

| **frequency (Hz)** | ***p* value (mean)** | ***p* value (Min)** | ***p* value (Max)** |
|---|---|---|---|
| **0.5** | 0.623 | 0.280 | 0.212 |
| **1** | 0.433 | 0.358 | 0.113 |
| **5** | 0.045 | 0.104 | 0.046 |
| **10** | 0.436 | 0.013 | 0.741 |
| **20** | 0.513 | 0.767 | 0.616 |
| **50** | 0.770 | 0.484 | 0.818 |

### Low-frequency ctACS induces phase-dependent changes in CS waveform

In contrast to the limited effects on the average CS waveform, ctACS induced significant phase-dependent changes at 0.5 Hz and 1 Hz stimulus frequencies (responses to higher frequencies will be treated separately below) in both the IS and SpkLt regions. This modulation is illustrated by responses of a PC to 0.5 Hz (Fig. 3a-c) and 1 Hz ctACS (Fig. 3g-i). The AAUC values for the IS (open black circles) and SpkLts (open red circles) parts of all CSs for each PC are plotted as a function of CS latency from the start of the AC cycle (Fig. 3a, g), and show the presence of a strong, phase-dependent modulation of the SpkLt waveform at both stimulus frequencies. For the SpkLt region, a sinusoidal fit to the AAUC data showed that the modulation was statistically significant in this PC (0.5 Hz: *A* = 1230; CI: 781-1679; 1 Hz: *A* = 1032; CI: 453-1611). For both frequencies, the normalized *A* (*A*/y_0_) was 0.241. For the IS region, only 0.5 Hz ctACS showed significant modulation (0.5 Hz: *A* = 118, CI: 45-191; 1 Hz: *A* = 83, CI: 51-115).

**Fig. 3.**
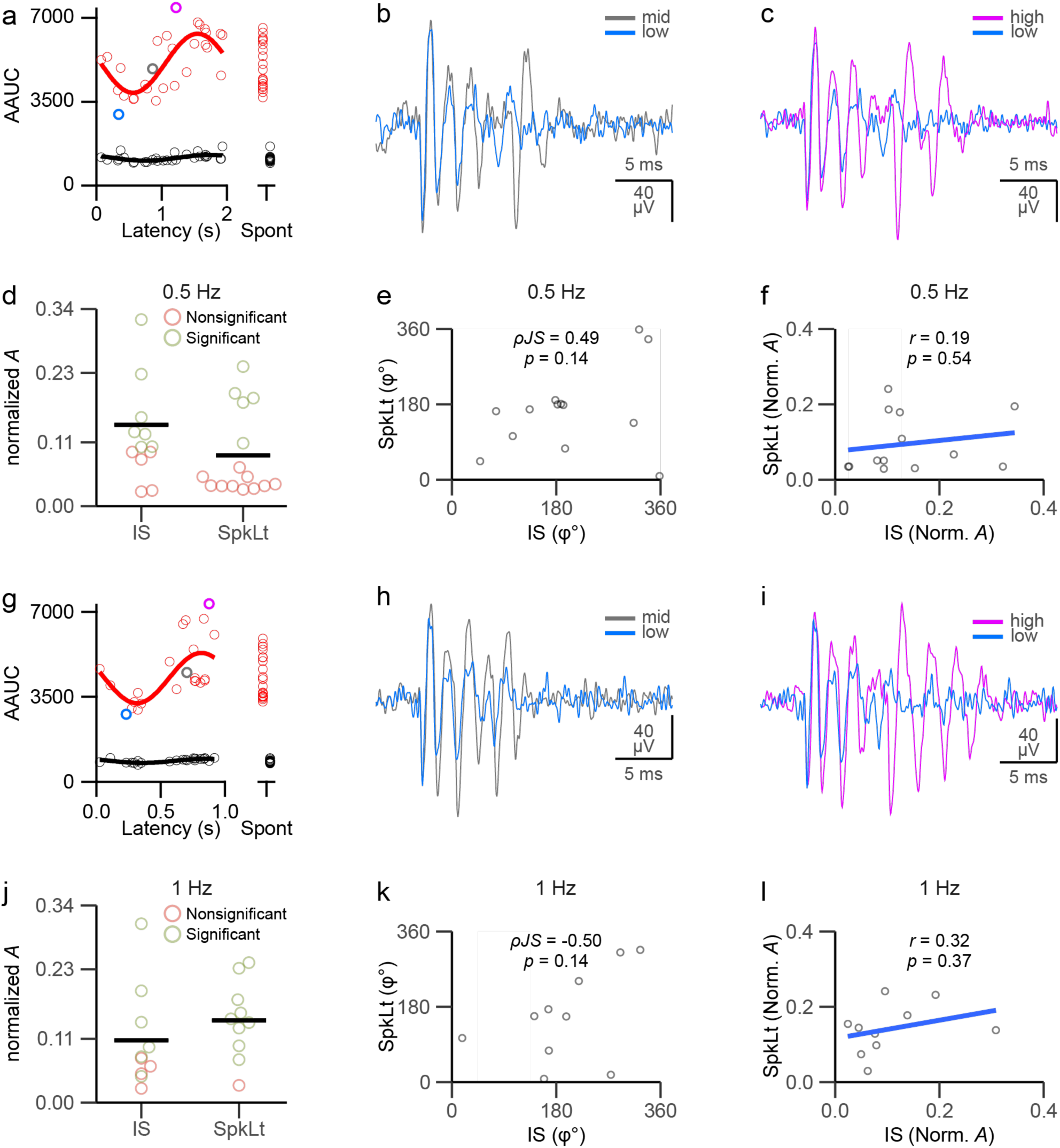
ctACS at 0.5 and 1 Hz induces phase-related changes in CS waveform. (a-f) Modulation during 0.5 Hz ctACS. **(a)** AAUC values during ctACS plotted as a function of the latency of the CS from the start of the AC cycle for all CSs of one PC: IS (open black circles) and SpkLt (open red circles). Solid curves show sine fits. AAUC values for spontaneous CSs (Spont) are plotted to the right to show the range of values normally present. **(b-c)** Example traces comparing CSs with small (blue) and moderate (gray) AAUCs **(b)** and those with moderate and large (purple) AAUCs **(c)**. Trace colors match the corresponding AAUC values shown in a. **(d)** Dot plot showing each PC’s normalized *A* (*A*/y_0_), with statistically significant values in green and nonsignificant ones in red for IS and SpkLts. **(e)** Scatterplot of phase of sine fit for IS versus SpkLt regions **(f)** Scatterplot of normalized *A* values for IS versus SpkLt regions. Blue is linear fit to the data. **(g-l)** Same as **(a-f)**, for CSs during 1 Hz ctACS.

The changes in CS waveform corresponding to these changes in AAUC can be seen in the CS traces, which compare CSs with low (blue trace), middle (grey trace), and high (purple trace) AAUC values (Fig. 3b-c, h-i, corresponding AAUC values indicated by matching color circles on panels 3a and 3g). Examination of these traces shows that changes in AAUC can reflect the addition/loss of spikelets and/or changes in spikelet amplitude and timing.

Across the population, both 0.5 Hz and 1.0 Hz ctACS produced phase-dependent changes in CS waveform; however, statistically significant modulation was found most often with 1.0 Hz ctACS. Panels 3d and 3j plot the normalized *A* for all PCs recorded during stimulation at these two frequencies. For the SpkLt period, 33.3% of the PCs (green circles, n = 5/15) demonstrated significant CS phase modulation at 0.5 Hz, whereas the percentage was 90% (n = 9/10) of PCs with 1 Hz ctACS. The average normalized *A* values at 0.5 Hz and 1 Hz were 0.088 ± 0.074 and 0.142 ± 0.066, respectively, again showing the somewhat stronger effect of the 1 Hz stimulation (Fig. 3d and 3j). However, considering only PCs that showed a statistically significant modulation, the normalized *A* values increase to 0.182 and 0.154. For the IS portion of the CS, the two stimulation frequencies were more similar in efficacy (0.5 Hz: n = 7/15 PCs; 1 Hz: n = 6/10 PCs). Overall, the normalized *A* values for the IS and SpkLt regions were not significantly different from each other at either 0.5 Hz or 1 Hz (0.5 Hz: paired t-test, *t*_(12)_ = 1.41, *p =* 0.18; 1 Hz: paired t-test, *t*_(9)_ =-1.21, *p =* 0.26). In sum, the data indicate that all parts of the CS can be altered by ctACS.

To assess whether changes in the IS and SpkLt regions of the CS were linked, we tested whether their phases (φ) and normalized *A* values from their sine fits were correlated. For both parameters, no significant correlation was found between the corresponding IS and SpkLt parameters for either 0.5 Hz (Fig. 3e; φ: *ρ_JS_* = 0.448, *p* = 0.141; Fig. 3f; normalized *A*: *r* = 0.19, *p* = 0.54) or 1 Hz (Fig. 3k; φ: *ρ_JS_* = -0.504, *p* = 0.136; Fig. 3l; normalized *A*: *r* = 0.32, *p* = 0.37) stimulation.

In sum, these findings show that low frequency ctACS can induce phase-dependent changes in CS waveform, and less often, changes in the average CS waveform, and that these changes reflect shifts within the range of the spontaneous CS waveforms rather than causing unphysiologically large or small CSs.

### ctACS modifies CS waveform over a wide range of ctACS frequencies

While measuring the effect on the entire SpkLt period of the CS gives information about overall phase-dependent effects of low frequency ctACS on that period, the SpkLt period can be quite extended in time. As a result, as the ctACS frequency increases, the SpkLt period can extend over a large fraction of the AC cycle, making assigning it a particular phase problematic. In addition, even at low ctACS frequencies, changes in CS waveform might be missed if countervailing changes occur in different parts of the SpkLt period that cancel each other’s effect on the AAUC. Thus, we subdivided the CS into smaller periods that roughly correspond to individual SpkLts for which a specific phase can still be meaningfully defined for higher ctACS frequencies. Analysis of these more restricted periods also allowed us to investigate whether changes in CS waveform were localized to specific parts of the CS.

We defined a SpkLt window for each detectable peak in the average CS and for estimated peaks on the tail end of the CS once detectable peaks could not be determined from the average (Fig. 1f, 4a; see Methods for details of the procedure for defining individual SpkLt windows). Using these smaller time windows, we found that modulation of the CS waveform was more common than suggested by examining the entire SpkLt window, and additional aspects about the modulation were revealed. The basic results are shown for one PC in figure 4b, where SpkLt windows 1, 2, and 4 had significant modulation to 10 Hz ctACS but the other windows did not (Win1: *A* = 314, CI: 238-389; Win 2: *A* = 401, CI: 297-506; Win 4: *A* = 209, CI: 399-19). For windows 1 and 2, the AAUC values follow the sine fit relatively closely, which suggests that the amplitude of the SpkLt in this window varies continuously during the AC cycle. The AAUC pattern for window 4 shows a different pattern, one which suggests an all-or-none type of modulation with phase. Specifically, the first half of the AC cycle primarily has small AAUC values, whereas during the second half of the cycle, both small and large AAUC values are present, with only a few intermediate values. These different distributions are reflected in the corresponding histograms of AAUC values in figure 4c. The difference in AAUC distributions for the two phases of the AC cycle for window 4 are most easily seen by plotting them separately (Fig. 4d). The positive phase has a unimodal distribution (bottom plot), whereas the negative phase has a bimodal distribution (top plot). For window 5, a bimodal distribution of AAUC values is even more evident (Fig. 4c). However, in this case, there was no phase-dependent modulation. The presence of both discrete and continuous variation in AAUC values suggests that ctACS modulates the CS waveform through two separable mechanisms: stepwise differences in the number of individual spikelets that make up the CS and graded changes in spikelet amplitude.

**Fig. 4.**
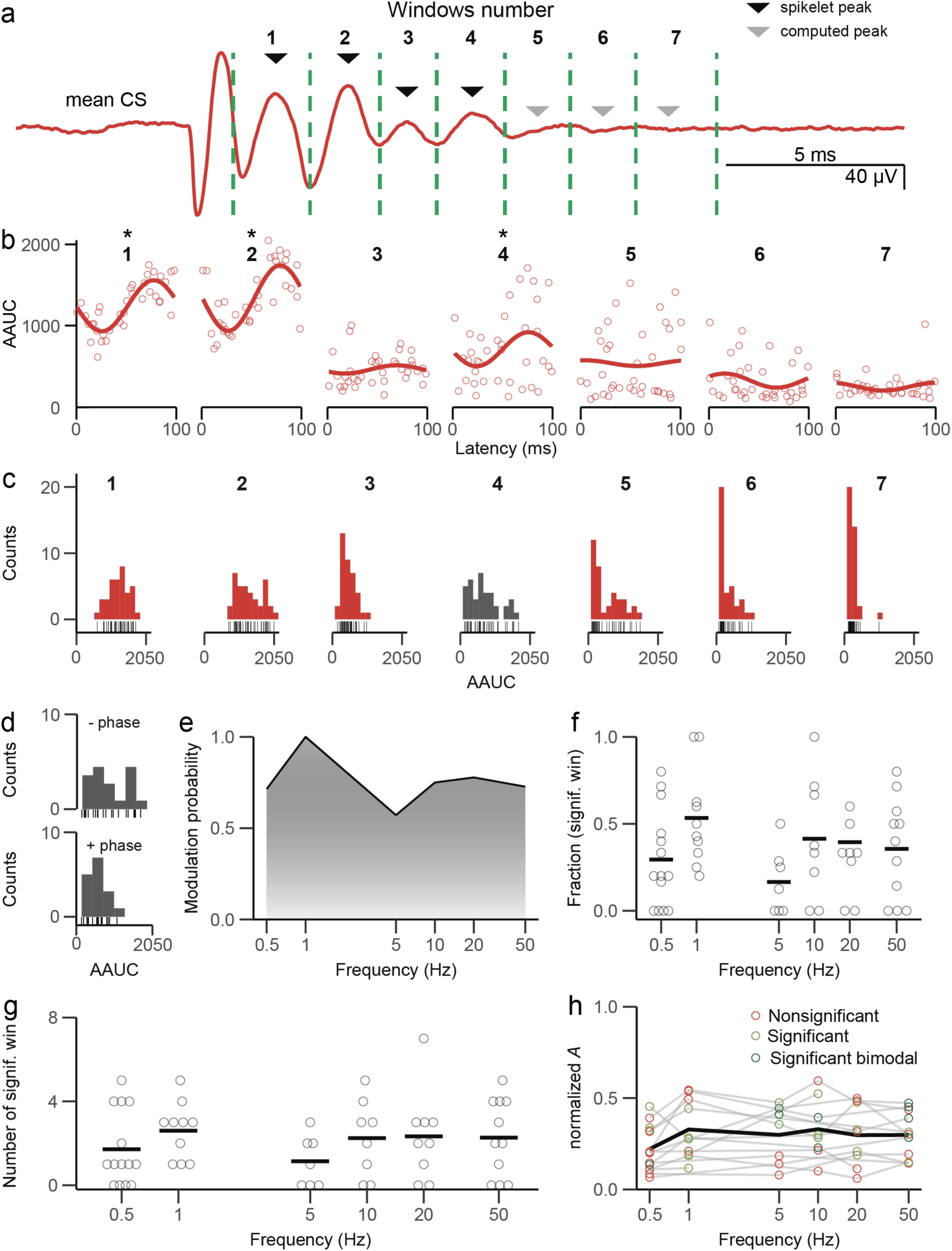
ctACS stimulation modulates individual spikelets with frequencies up to 50 Hz. **(a)** Mean CS waveform from one PC during 10 Hz ctACS. Four SpkLt peaks were detected from the mean waveform (black triangles) and three were computed (gray triangles). Green dashed lines indicate window limits surrounding each peak. **(b)** AAUC values for each window in **(a)** plotted as a function of the latency of the window from the start of the AC cycle. Sine fits are indicated by solid curves. **(c)** Histograms of AAUC for each window for PC in **b**. The dark grey histogram (window 4) shows phase modulation and evidence for present or absent spikelets (bimodal histogram), while window 5 shows a clear bimodal pattern. **(d)** Data in window 4 of panel **c** replotted with CSs during positive (0-180°; 0 - 50 ms) and negative (180-360°; 50 - 100 ms) phases of AC cycle plotted separately. **(e)** Probability of significant modulation in at least one spikelet window across stimulation frequency. **(f)** Overall probability of a window showing significant modulation for each PC at each ctACS frequency. Horizontal bars indicate the population means. **(g)** Number of significant windows for each PC at each frequency. Horizontal bars indicate the population means. **(h)** For each PC, the window with the most extensive modulation (highest normalized *A*) was plotted for each frequency. The black line indicates the mean at each frequency.

Across the population, almost all PCs (n = 15/16) showed significant AAUC modulation in at least one individual SpkLt window during ctACS in at least one tested frequency (Fig. 4e). A majority of PCs (n = 9/16) had significant modulation in at least one window across all tested frequencies. The fraction and absolute number of windows with significant modulation in each cell are shown in panels 4f and 4g, respectively, and did not vary systematically with frequency. On average, significant phase modulation occurred in 2.05 windows per PC, or 37.7% (n = 122/324) of all windows taken from all PCs and ctACS frequencies. To test whether modulation strength varied with ctACS frequency, the window with the highest normalized *A* for each PC was used to determine the maximal effect of ctACS on the CS waveform (Fig. 4h). Averaging across all frequencies, the mean maximal normalized *A* was 0.289 ± 0.147. This value did not show any linear trend with frequency (LMM: normalized *A* ∼ stimulus frequency; *F*_(1, 45.3)_ = 0.24, *p =* 0.63). It is worth noting that this lack of frequency dependence contrasts with the strong dependence of SS firing rate modulation by ctACS and raises the possibility of being able to shift the balance of ctACS’s action between altering CS waveform and modulating SS output by using different ctACS frequencies.

In sum, ctACS appears to cause both discrete phase-dependent jumps in the AAUC that could reflect the addition or subtraction of individual spikelets and/or more continuous changes in the AAUC that could reflect alterations in SpkLt amplitude in most PCs.

### ctACS-phase-induced changes in AAUC are correlated with spikelet amplitude and number

The above results suggest that ctACS-induced changes in the AAUC could reflect changes in the number and/or the amplitude of the CS spikelets. To begin to determine the extent to which each of these spikelet parameters is affected by ctACS, the AAUC histograms for the SpkLt windows were categorized as either unior bi-modal by visual inspection. Of the 122 windows that showed a significant AAUC phase modulation during ctACS, 22.95% (n = 28/122) exhibited a bimodal distribution, suggesting a discrete jump in the CS waveform AAUC. Moreover, at least one such window was found in a majority of PCs (62.5%, n = 10/16). It is worth noting that this percentage may be an underestimate of bimodal AAUC distributions to ctACS, as some PCs with strong modulatory characteristics may have been classified as nonsignificant because of increased variance due to the discrete distribution. Consistent with this possibility, a number of windows with large normalized *A* values (≥ 0.2) that were not statistically significant showed a bimodal distribution (26/324). If these 26 cases are added to the total of bimodal histograms of cells with significant modulation, the percentage rises to 36% (54/148) of windows with evidence ctACS inducing and blocking spikelets.

To investigate further the contribution of spikelet number and amplitude to changes in AAUC induced by ctACS, we directly measured these two parameters for a subset of PCs with high signal to noise recordings, which allowed individual SpkLts to be counted at frequencies of 0.5 Hz (n = 15) and 1 Hz (n = 10). These frequencies were chosen because CS duration is relatively short compared to the cycle periods at these frequencies, allowing assignment of a specific phase to the CS.

We first assessed the relationship between AAUC and the number and/or amplitude of the spikelets during both ctACS and spontaneous activity. AAUC was correlated with both SpkLt number and average amplitude during ctACS at 1 Hz and spontaneous activity (Fig 5a, b, d, and e). Across the PC population, AAUC and SpkLt number were significantly positively correlated during ctACS and during spontaneous activity for most PCs (spontaneous: 9/10; 0.5 Hz ctACS: 12/15 PCs; 1 Hz ctACS: 6/10 PCs; Fig. 5c). Similar results were found for AAUC versus SpkLt amplitude (spontaneous: 7/10; 0.5 Hz; n = 10/15, 1 Hz; n = 9/10, Fig. 5f).

**Fig. 5.**
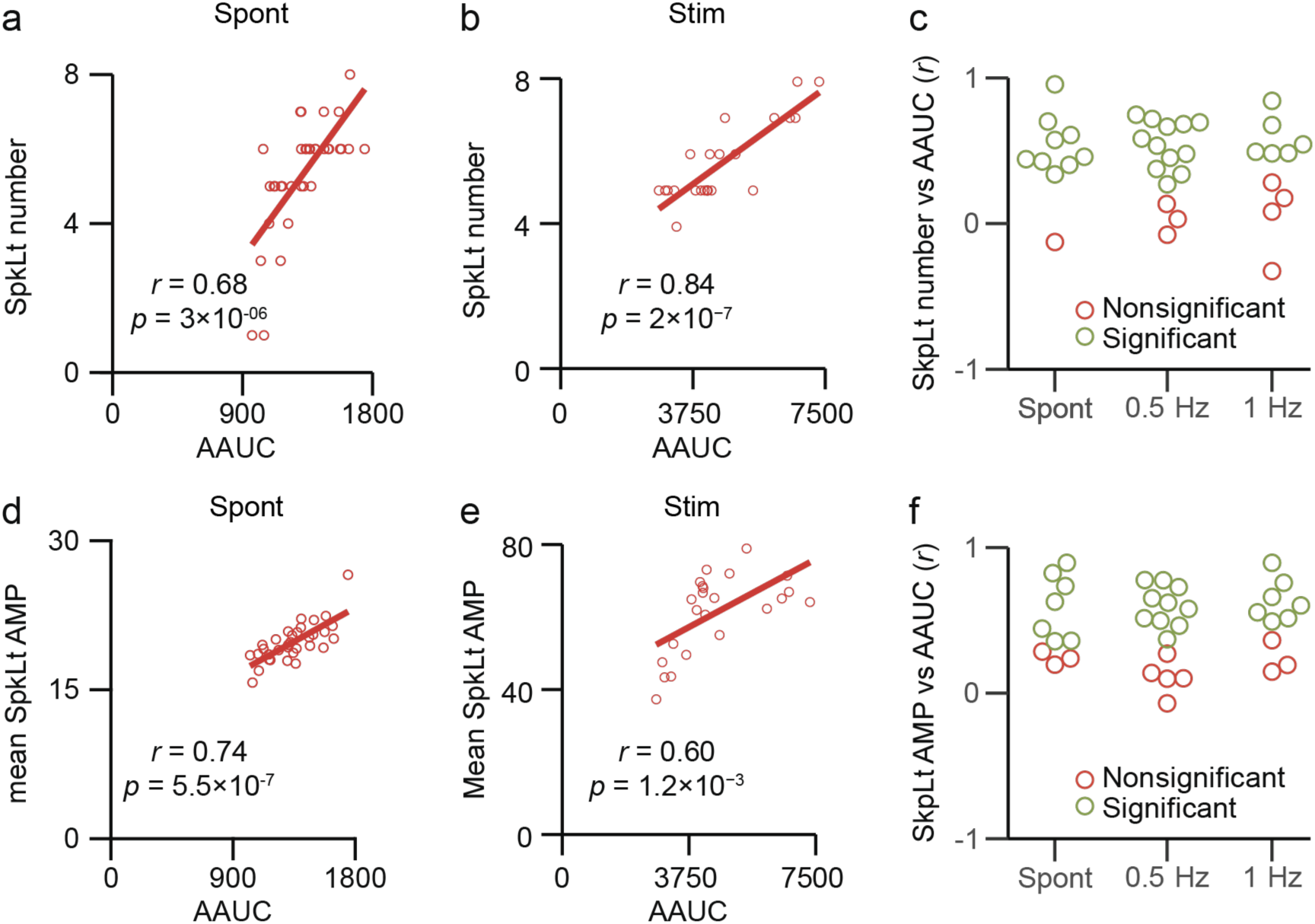
Spikelet number and amplitude correlate with AAUC. **(a)** Correlation of AAUC and SpkLt number for spontaneous CSs from one PC. **(b)** Same as **a** for CSs during ctACS. **(c)** Correlation of AAUC and SpkLt number for all PCs for spontaneous (Spont) activity and during ctACS. Green open circles indicate a significant correlations (r ≠ 0, *p* < 0.05). **(d)** Correlation of AAUC and SpkLt amplitude for spontaneous CSs from a different PC. **(e)** Same as **d** for CSs during ctACS. **(f)** Correlation of AAUC and SpkLt amplitude for all PCs for spontaneous (Spont) activity and during ctACS.

Thus, changes in both SpkLt number and amplitude correlate with changes in AAUC as expected. This suggests that CS modulation, as measured by AAUC, likely reflects changes in SpkLt number and amplitude, raising the probability that SpkLt number and amplitude are modulated by ctACS.

To determine whether ctACS causes a change in the average SpkLt numbers or amplitude from spontaneous CS activity, these parameters were compared between the ctACS and spontaneous activity periods. For both SpkLt number and amplitude, 1 Hz ctACS caused a significant change in the mean number and amplitude of SpkLts in only 4 of 10 PCs (different set of PCs for each parameter; Fig. 6a, b). Similar results (number: 4 of 15; amplitude 2 of 15 PCs) were found for 0.5 Hz ctACS (data not shown). Overall, the percent changes between ctACS and spontaneous values were small (0.5 Hz: number, 10.0 ± 12.7%; amplitude, 5.64 ± 5.04%; 1.0 Hz: number, 9.14 ± 10.2%; amplitude, 6.27 ± 5.56%). There were no significant change in SpkLt number and amplitude over the whole population: (0.5 Hz: number, *t*_(12)_ = 0.139, *p =* 0.221; 1 Hz: number, *t*_(9)_ = 1.06, *p =* 0.319) and (0.5 Hz: amplitude, *t*_(12)_ = 0.386, *p =* 0.707; 1 Hz: amplitude, *t*_(9)_ = 0.032, *p =* 0.975). In sum, the relatively small changes in spikelet parameters between ctACS and spontaneous periods are consistent with the AAUC results.

**Fig. 6.**
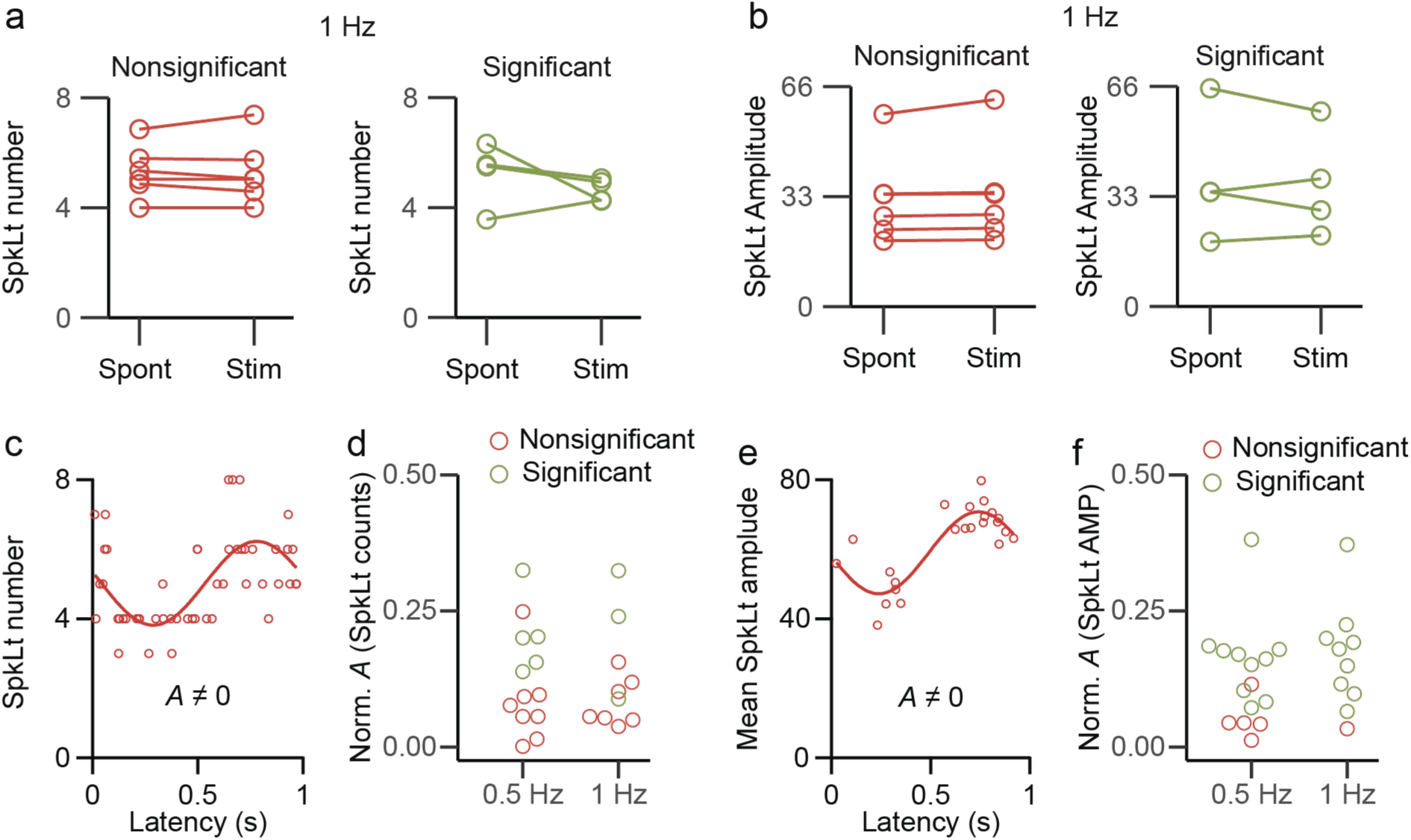
ctACS phase-related modulation of SpkLt parameters. **(a)** Pair plot comparing the mean spikelet number during spontaneous activity (Spont) and stimulation (Stim) grouped by significance of difference in the two conditions. **(b)** Same as **a** but for mean spikelet amplitude. **(c)** Modulation of SpkLt number during AC cycle for CSs from one PC. N **(d)** Plot of normalized *A* value for each PC for sine fits to spikelet number versus phase, at 0.5 Hz and 1 Hz. Green open circles indicate significant modulation; red open circles indicate nonsignificant modulation. **(e-f)** Same as **d-e**, for sine fits to spikelet amplitude versus phase.

Next, we investigated whether SpkLt number and amplitude showed phase-related variations during ctACS. Figure 6 shows an example of a PC with significant modulation in SpkLt number (Fig. 6c; *A* = 1.21, CI: 0.77-1.64; normalized *A* = 0.24) and another PC with significant modulation of SpkLt amplitude (Fig. 6e; *A* = 11.78, CI: 8.8-14.7; normalized *A* = 0.200). Across the PC population, SpkLt amplitude (normalized *A*) was significantly modulated in most PCs (0.5 Hz: 10/15; 1 Hz: 9/10; Fig. 6f). Only a minority of PCs showed a statistically significant modulation in SpkLt number (0.5 Hz: 4/15, 1 Hz: 2/10; Fig. 6c). The greater number of PCs with significant SpkLt amplitude modulation than SpkLt number modulation could suggest that the former is the major effect of ctACS on CS waveform. However, a possible reason for the lower number of significant PCs for SpkLt number is the discrete nature of that parameter causing an increase in variance. The effect of this can be seen by comparing the distribution of data points about the fitted sine wave for the two examples in figures 6c and 6e. Note the small scatter about the sine wave for the SpkLt amplitude data, which contrasts with the wider dispersion about the sine wave for the SpkLt number data, because of the discrete jumps in spikelet number. This increased variability could lead to cases where a relatively large change in normalized *A* values with phase is still not statistically significant.

In sum, both SpkLt parameters can be modulated by ctACS and contribute to the changes in AAUC described earlier. The evidence is strongest for SpkLt amplitude being affected, but there is also evidence for SpkLt number being affected in at least some PCs, and it is possible that our methodology underestimates the true effect of ctACS on SpkLt number.

### SS modulation correlates with changes in the CS waveform

We previously showed that SSs are significantly modulated in a frequency-dependent manner during ctACS (Mourra et al., 2025). The strength of this modulation should reflect the effective electric field that the PC experiences during ctACS and thus could be used to help quantify the relationship of the electric field to changes in CS waveform induced by ctACS. The above results show that 1 Hz ctACS was most consistent in modulating CS waveform. Therefore, we examined ctACS-induced modulation of SS activity at 1 Hz, which had not been previously assessed. Consistent with our prior results at other stimulus frequencies, ctACS did not alter the average SS firing rate (spontaneous, 31.5 ± 19.7 Hz; ctACS, 30.5 ± 19.4 Hz; *t*_(9)_ = 1.54, *p =* 0.158) but did produce a phase-locked modulation of SS activity in PCs (Fig. 7a, b). The strength of the SS modulation was quantified by the magnitude of the CMV (r̅ = 0.283 for the example PC shown in Figure 7c). At 0.5 Hz, the PC population had a mean r̅ of 0.182 ± 0.219, and most PCs (13/15, 86.7%) showed significant SS modulation. At 1 Hz, the PC population had a mean r̅ of 0.323 ± 0.201, and all PCs (10/10, 100%) showed significant SS modulation. Even stronger modulation was seen at higher stimulation frequencies (50 Hz: 0.403 ± 0.287; 80 Hz: 0.453 ± 0.293; LMM: SS r̅ ∼ frequency; *F*_(6, 81)_ = 12.904, *p* = 3.4 × 10^-10^), and the Tukey-adjusted post hoc comparison showed that modulation strength in response to 1 Hz stimulation was significantly lower than in response to 50 and 80 Hz stimulation (1 Hz vs 50 Hz, *t*_(85)_ = -3.79, *p* = .005; 1 Hz vs 80 Hz, *t*_(84)_ = -5.73, *p* = 3.17×10^−6^). Consistent with previous reports (Avlar et al., 2023; Mourra et al., 2025), the individual t̅’s (phase of the CMV) clustered around either 90° or 270°, with the large majority of PCs having a t̅ close to 270° (Fig. 7d, e).

**Fig. 7.**
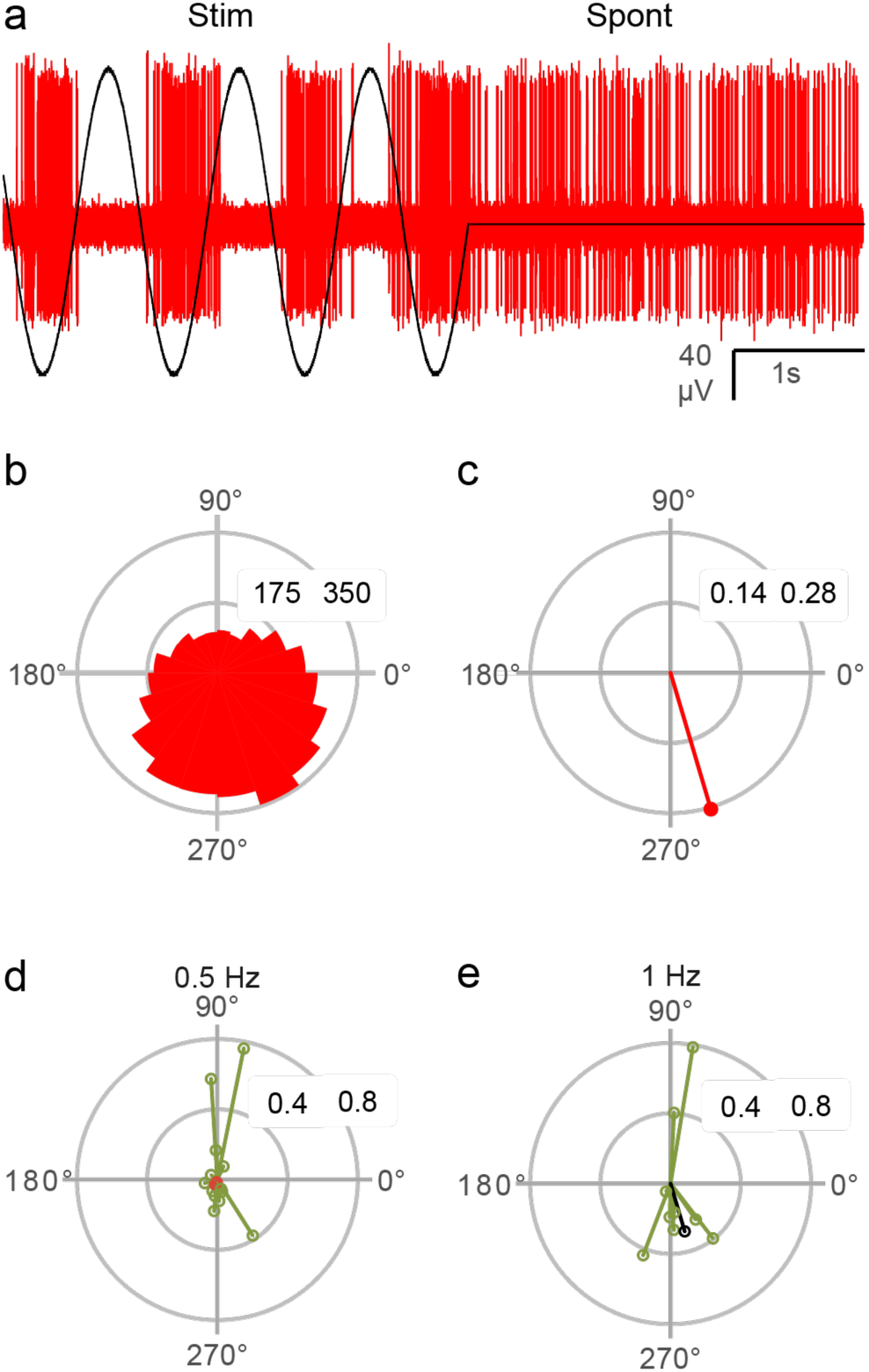
ctACS-induced modulation of SS firing rate. **(a)** Example of extracellular recording from a PC (red trace) showing significant response to 1 Hz ctACS and spontaneous activity. Black trace indicates ctACS stimulation period. **(b)** Polar histogram of the timing of SS activity relative to the start of each sinusoidal cycle. Bins set to 1/20 of the cycle period; the radial axis shows spike count. **(c)** Polar plot of the circular mean vector for the PC spike activity from **b**. **(d)** Polar plot showing the phase and magnitude of each PC’s circular mean vector for activity during 0.5 Hz ctACS; green indicates significant modulation, while red indicates nonsignificant modulation. **(e)** Same as **d** for 1 Hz ctACS. The black vector corresponds to the PC in **a-c**. Green indicates significant modulation.

To facilitate comparison of the phase relationship of the SS and CS modulations, we added 90° to the phase of the sine fit to the CS AAUC values (φ* = φ + 90°), which allowed a direct comparison of the phase of the peak of the sine wave fit to the CS data to the peak of SS activity given by the direction of the CMV (t̅). This analysis was limited to PCs for which both the CS AAUC and SS activity were significantly modulated during either 0.5 and 1 Hz ctACS (n = 10 PCs; for PCs that were tested and modulated at both frequencies, the frequency with the stronger SS modulation was used (n = 4 PCs)). The t̅ and φ* for these PCs are plotted in figures 8a and 8B, respectively. The SS t̅ values clustered around 270° (circular mean = 276°, n = 7 PCs) and 90° (circular mean = 80°, n = 3 PCs), forming a bimodal distribution. In contrast, the φ* phases for CS AAUC modulation were more widely distributed (uniform distribution, Rayleigh test). We then investigated the relationship between the t̅ and φ* values of each PC. For each cluster of t̅ values, the t̅ and corresponding φ* values are connected in plots of figure 8c. While some PCs (n = 3) showed close to a 180° difference, most did not.

**Fig. 8.**
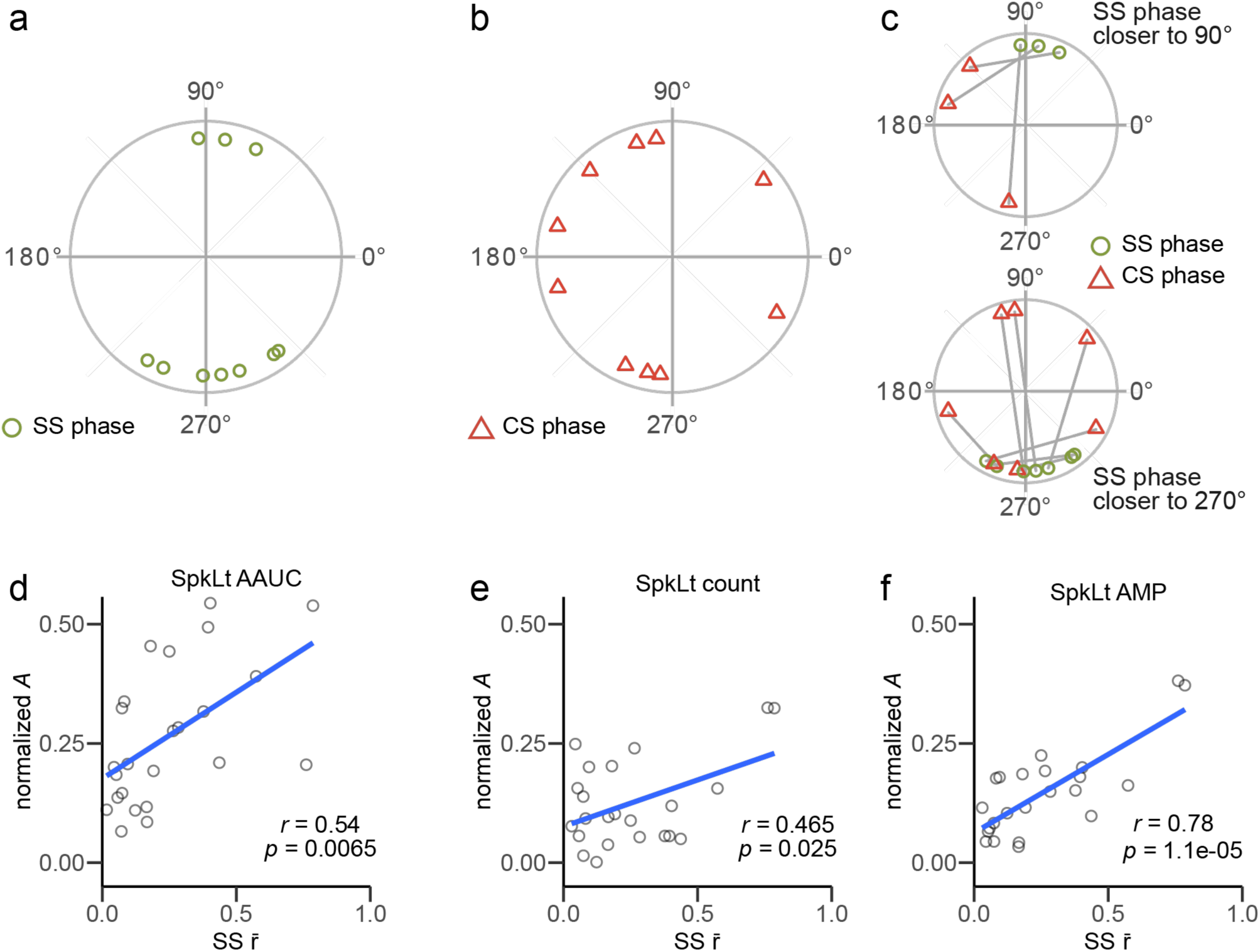
CS waveform modulation correlates with SS firing rate changes. **(a)** A polar plot showing the phase of the SS mean circular vector (t̅) of each PC. **(b)** A polar plot showing the φ* (phase of the sine fit plus 90°) for the CS AAUC modulation. **(c**) Data in **a** and **b** overlaid with a gray line connecting the t̅ and φ* of each PC. Top plot shows PCs with t̅’s close to 90° and bottom plot shows PCs with t̅’s closer to 270°. **(d-f)** Relationship of SS r̅ to modulation (normalized *A*) of AAUC **(d)**, SpkLt count **(e)**, and SpkLt amplitude **(f)** during 0.5 and 1 Hz ctACS.

Next, we examined the relationship between SS r̅ and the parameters that measured the amplitude of the CS waveform modulation: normalized *A* for the windows with highest value for the PC for the AAUC, SpkLt number, and SpkLt amplitude parameters. The results were collapsed over 0.5 - 1 Hz (n = 24; 10 PCs at both frequencies, 4 at 0.5 Hz only). Notably, there was a significant positive correlation between SS r̅ and the strength of CS waveform modulation across all measured parameters (Fig. 8d-f).

In sum, changes in CS waveform induced by ctACS are correlated with its modulation of SS activity. Assuming SS modulation reflects the strength of the effective electric field induced by ctACS implies that the magnitude of the CS changes is proportional to the field strength.

### ctACS-induced modulation depends on recording depth

Our PCs were located at different depths, and the strength of the electric field generally, but not always, decreases with distance (Asan et al., 2019b; Avlar et al., 2023). Thus, we investigated how the effect of ctACS varied with the depth of the PC for both the CS waveform modulation and the SS firing rate modulation at each stimulus frequency. SS r̅ and recording depth were significantly negatively correlated at 0.5, 5, 20, and 50 Hz (at 1 and 10 Hz the correlation was negative but not significant), with the strongest correlations occurring at the higher frequencies (Fig. 9a).

**Fig. 9.**
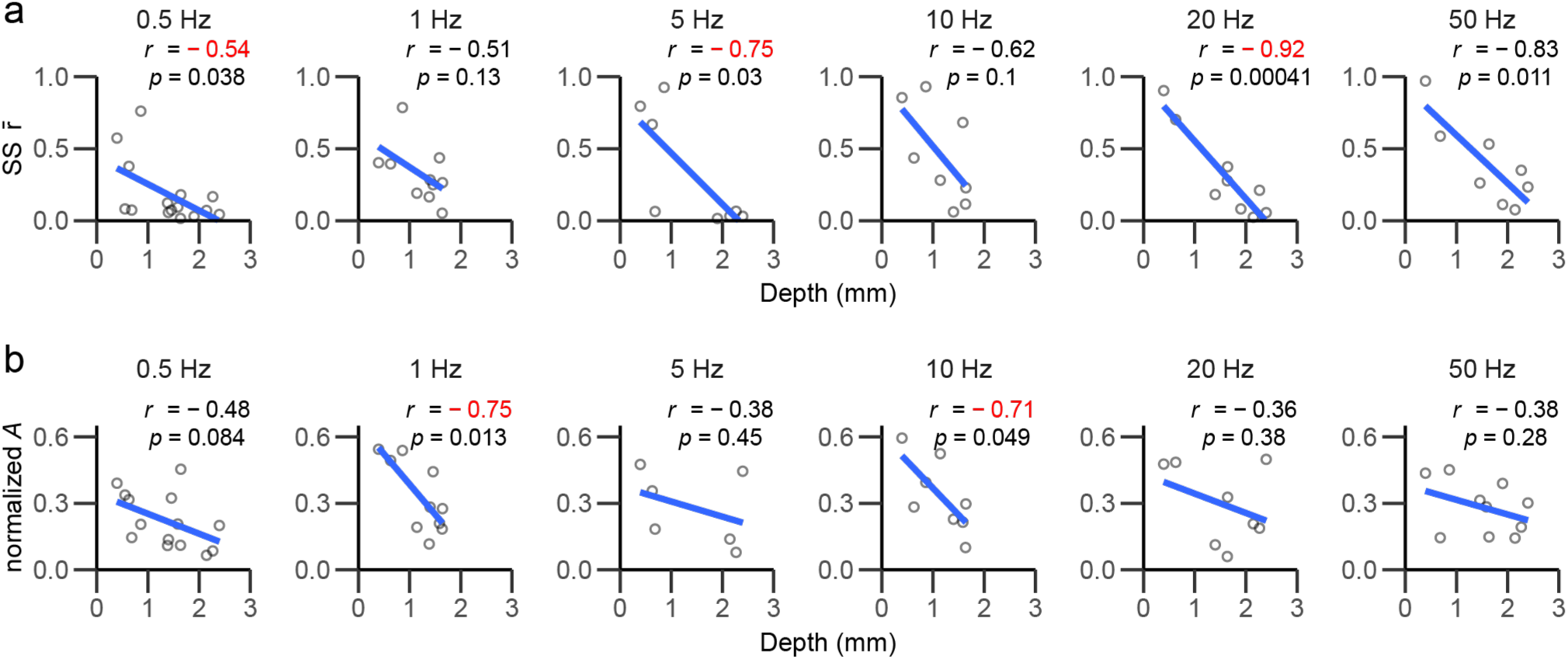
ctACS-induced waveform modulation and SS entrainment vary with recording depth. **(a)** SS r̅ plotted against recording depth for each stimulus frequency. Each data point is from one PC; the blue line indicates the best-fit line at each frequency tested. **(b)** Normalized *A* plotted against recording depth for each stimulus frequency; the blue line indicates the best-fit line at each frequency tested. Normalized *A* is calculated using the AAUC for the window with the highest normalized *A* for each PC (see Fig. 4h).

To evaluate the relationship between CS waveform modulation and recording depth at each frequency, we used the window with the highest normalized *A* value for each PC (Fig. 4h). Again, we found the effect of ctACS decreased with recording depth, although this relationship was less consistent overall than that for SS modulation (Fig. 9b), being significant only at 1 and 10 Hz (trending at 0.5 Hz). The contrast with SS entrainment is worth noting: for CS waveform, the correlation was strongest at lower frequencies (0.5 - 10 Hz).

To probe further the relationship between PC depth and the frequency dependence of the CS waveform and SS firing-rate modulation, we assessed how modulation strength depended on the three variables: stimulus frequency, PC depth (continuous), and modulation type (discrete). We found a significant 3-way interaction (LMM: modulation strength ∼ stimulus frequency * PC depth * modulation type; *F*_(1, 89)_ = 4.87, *p* = .03) in addition to main effects for all three variables (frequency, *F*_(1, 92)_ = 7.24, *p* = 0.008; PC depth, *F*_(1, 16)_ = 8.26, *p* = 0.01; modulation type, *F*_(1, 89)_ = 4.42, *p* = 0.036) and significant 2-way interactions (frequency* modulation type, *F*_(1, 89)_ = 11.8, *p* = 0.0008; PC depth* modulation type, *F*_(1, 89)_ = 6.30, *p* = 0.013).

To better visualize the 3-way interaction, PCs were grouped based on their depth relative to the cerebellar surface (0 - 1mm, 1 - 2 mm, and 2 - 3 mm), and the strength of the ctACS induced modulation was plotted as a function of stimulus frequency (Fig. 10). The greatest divergence between CS waveform and SS modulation occurred in PCs located at more superficial depths (0-1 mm) from the cerebellar surface. Specifically, mean normalized *A* values (red trace) for CS waveform modulation did not vary across frequencies at any depth, while mean SS r̅ values (green trace) increased with frequency at more superficial depths; the normalized A and SS r̅ were less divergent at deeper PC depths.

**Fig. 10.**
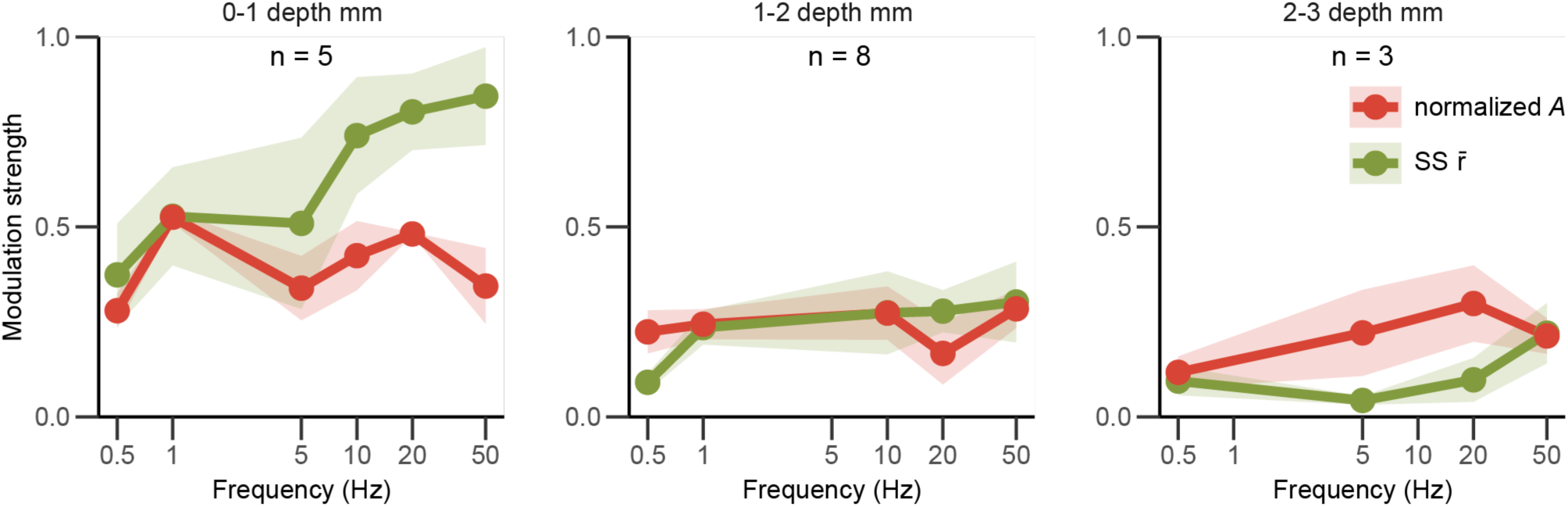
CS waveform and SS modulation divergence across frequency and depth. The mean normalized *A* and SS r̅ are plotted as a function of frequency for PCs grouped by recording depth: 0-1 mm, 1-2 mm, and 2-3 mm from the cerebellar surface. Normalized *A* is calculated using the AAUC for the window with the highest normalized *A* for each PC (see Fig. 4h). Shaded regions show one SEM.

In sum, these results indicate that the strength of ctACS-induced modulation of PC activity decreases with dorsoventral distance from the stimulation electrode on the dorsal skull, and that SS modulation is highly frequency dependent for superficially located PCs, whereas CS modulation did not differ significantly by stimulation frequency.

## Discussion

In this paper, we investigated how ctACS alters CS firing rate and waveform *in vivo*. Changes in CS waveform were quantified in multiple ways (AAUC and SpkLt numbers and amplitudes), all of which showed that ctACS induces phase-dependent changes in CS waveform. To our knowledge, these results are the first to show that ctACS induces a phase-dependent modulation of the CS waveform *in vivo*. As discussed below, such changes in CS waveform could have both short and long term effects on the cerebellar cortex and possibly the cerebellar nuclei: effects that could underlie ctACS’s ability to cause sustained changes in brain function and contribute to ctACS’s therapeutic actions. Modulation of CS firing rate was less frequently observed; however, it too may represent a way to modify synaptic plasticity and target ctACS’s actions.

### ctACS modulates CS waveform by altering spikelet numbers and amplitudes

The strongest and most consistent effect of ctACS on CS activity was to induce phase-dependent changes in CS waveform. In contrast, relatively little change in the average CS waveform from spontaneous CSs was observed, likely due to the symmetric nature of ctACS. Analysis of AAUC changes showed that both the IS and the following SpkLts portions of the CS were significantly modulated by ctACS. The modulation was strongest in PCs located on or near the brain surface (apices of the folia) but was also present in PCs along folial walls. When AAUC values were calculated for specific, smaller, discrete time windows that roughly corresponded to individual spikelets, a large majority of PCs showed significant modulation at each of the tested frequencies, with 1 Hz stimulation being the most effective in terms of the percentage of modulated PCs; however, modulation strength was similar across stimulation frequencies.

ctACS did not cause a generalized change in the overall CS waveform but rather caused modulation of discrete parts of the CS, corresponding to specific spikelets, as shown by the analysis of individual SpkLt windows (e.g., Fig. 4, in which only three of seven windows showed significant modulation). Confirming these AAUC window analyses, direct measurement of spikelet parameters showed that ctACS caused changes in both their numbers and amplitudes. These changes likely explain much of the overall effect on the AAUC, as they correlated highly with AAUC values (Fig. 5).

It is worth noting that the AAUC and SpkLt analyses each have limitations and strengths and thus complement each other. SpkLt analyses provide more direct measures of specific parameters than do the AAUC analyses but are limited in that they could be performed only on a subset of PC recordings with particularly high signal-to-noise ratios (SNR) that allowed clear identification of individual spikelets. Moreover, even in such cases, defining the exact cutoff between ‘true’ spikelets and fluctuations in baseline noise can be difficult. In contrast, AAUC analyses provide only general information about waveform changes, but they avoid the necessity of choosing arbitrary cutoffs. The consistency between these two approaches suggests that the findings are robust.

Our data do not definitively identify the mechanisms by which ctACS induces changes in CS waveform. However, our data do suggest multiple interacting factors are involved. Applied electric fields polarize neurons directly, modifying spiking, and in the case of the PC specifically, evoke calcium spikes in PC dendrites *in vitro* that could be seen at the soma (Chan et al., 1988). Such spikes would alter the form of the CS. Since direct polarization is maximal at the peak and trough of the AC cycle, the largest changes in CS waveform should have occurred at these limits of the AC cycle, if direct polarization of the PC membrane is driving changes in CS waveform. For a slim majority of our PCs, this was indeed the case. However, this pattern was far less consistently observed than it has been for SS modulation (i.e., maximal SS firing rates generally occur close to in-phase or 180° out-of-phase with respect to the ctACS cycle (Mourra et al., 2025). Moreover, *in vitro* Ca spikes were evoked 180° out-of-phase with SS activity, whereas here both in- and 180° out-of-phase relationships were observed, as well as other phase relationships (Fig. 8). The distribution of CS waveform modulation phase (i.e., φ*) across the PC population suggests that additional factors beyond direct PC membrane polarization are at least partly responsible for this modulation. One likely candidate is modulation of molecular layer interneurons, as their activity can alter CS waveform (Eccles et al., 1966; Rowan et al., 2018) and can be modulated by applied electric fields (Chan and Nicholson, 1986). Another possibility is that ctACS may affect the state of IO neurons (most likely indirectly, via modulation of SS activity and thereby modulation of cerebellar nuclear neurons that project to the IO), as the state of the IO affects the number of spikes comprising each IO neuronal discharge, which in turn affects the number of CS spikelets (Maruta et al., 2007; Mathy et al., 2009; Bazzigaluppi et al., 2012; De Gruijl et al., 2012). However, the general lack of CS firing rate modulation makes this a less likely possibility (see below).

In sum, the CS waveform modulation of about half of our PC population may largely be explainable by a direct polarization of the PC membrane by the electric field generated by the ctACS. For the other PCs, alternative mechanisms likely play a significant role. In these cases, it may be that the PC experiences a weaker effective field because of its orientation, allowing the synaptic effects of interneurons or other processes to become dominant. Further experiments with synaptic blockers could test these possibilities.

### ctACS modulation of CS waveform may have short- and long-term effects on cerebellar activity

Studies have shown that CSs can have both short-term and long-term effects on cerebellar cortical activity. On the short time scale, CSs can induce a number of changes in cerebellar cortical activity. These include the well-known post-CS pause in SS activity, as well as post-pause modulation of SS activity for hundreds of milliseconds (Granit and Phillips, 1956; Bell and Grimm, 1969; Murphy and Sabah, 1970; Bloedel and Roberts, 1971; Latham and Paul, 1971; Burg and Rubia, 1972; Ebner and Bloedel, 1981a; Ebner and Bloedel, 1981b; McDevitt et al., 1982; Sato et al., 1992). Changes in CS firing rates can also lead to inverse changes in SS firing rates (Marshall and Lang, 2009; Chaumont et al., 2013), which could be consistent with a homeostatic role for CS activity. On a longer timescale, CSs have long been thought to be required for plasticity at the PF to PC synapse, most notably LTD (Ito, 2001).

Changes in CS waveform are likely an important parameter of these short and long term effects on cortical activity, and most specifically, PC activity. Early reports on the relationship of SpkLt numbers and short-term SS levels are mixed, with one study finding a correlation (Gilbert, 1976) and another failing to do so (Mano, 1970). However, a more recent study found correlations between SpkLt numbers and SS firing rate both before and after the CS (Burroughs et al., 2017). Of potentially more relevance for using ctACS for long-term changes in cerebellar function is that variations in SpkLt number have been shown to affect the type of plasticity induced by the CS in several different paradigms and preparations (Mathy et al., 2009; Rasmussen et al., 2013; Yang and Lisberger, 2014; Rowan et al., 2018) but see (Titley et al., 2019). In these studies changes of 1-2 spikelets altered the strength of the plasticity or reversed its polarity (Mathy et al., 2009; Rasmussen et al., 2013). This raises the issue of whether ctACS’s effect on spikelet number is large enough to affect plasticity. The average normalized *A* value for our PC population was 0.128 for 0.5 Hz stimulation and 0.123 for 1 Hz stimulation (Fig. 6d), corresponding to peak-to-peak modulations of 0.256 and 0.246, respectively. Given that the average number of SpkLts/CS was 6.19 (0.5 Hz) and 5.28 (1 Hz), the average peak-to-peak variation in SpkLt number is 1.58 for 0.5 Hz and 1.30 for 1 Hz across the AC cycle, suggesting that ctACS should be able to influence the strength and polarity of CS-induced plasticity significantly.

The above discussion focuses on changes in SpkLt numbers as a possible mechanism for ctACS to alter plasticity. However, in addition to changes in SpkLt number, changes in spikelet amplitude were observed, and in fact, these changes were found most often. The relationship of SpkLt amplitude to PC plasticity has not been investigated to our knowledge; however, it is very plausible that variations in spikelet amplitude also could have a significant effect.

Beyond effects on cerebellar cortical plasticity, variations in SpkLt number may affect cerebellar output. Both *in vivo* and *in vitro* studies show that a CS may result in more than one spike propagating down the PC axon (Ito and Simpson, 1971; Khaliq and Raman, 2005; Monsivais et al., 2005). Moreover, high frequency (500 Hz) spike bursts, like those comprising the CS, may have a distinct, long lasting effect on the state of the postsynaptic cerebellar nuclear neurons, shifting them from an excited spiking state to a silent one (see figure 3 of Jahnsen, 1986). The phase-dependent variations in the total number of SpkLts by ctACS found here do not directly show that the number of spikes propagated down the PC axon is affected by ctACS, and indeed, variations in the total number of SpkLts associated with changes in CS synchrony may occur without changes in the number of presumed axonally-propagated spikes (Lang et al., 2014). However, under *in vitro* conditions, SpkLt amplitude has been shown to be a marker for whether it is associated with an axonally propagated spike (i.e., larger amplitude spikelets have a higher probability of being propagated) (Khaliq and Raman, 2005; Monsivais et al., 2005). Thus, the fact that we found significant modulation of SpkLt amplitude by ctACS in a majority of PCs suggests that ctACS may alter the number of propagated axonal spikes generated by a CS.

Collectively, the effects on CS waveform described here suggest that ctACS may be able to modify synaptic plasticity in the cerebellar cortex and the effect of PC activity on the cerebellar nuclei. Both of these actions may contribute to ctACS’s ability to cause long-lasting changes in brain function in addition to altering the ongoing cerebellar output.

### ctACS modulation of CS firing rate

In contrast to the prevalence of CS waveform modulation, significant modulation of CS firing rate was observed infrequently. Only one or zero PCs showed CS firing rate modulation at the highest (20, 50 Hz) and lowest (0.5, 1 Hz) stimulus frequencies, respectively. At intermediate frequencies (5, 10 Hz), a somewhat higher percentage, but still a minority, of PCs (37.5%; 3/8 PCs in each case) were modulated. This was a bit surprising given the strong modulation of SS firing rate and that changes in SS levels are known to cause changes in CS activity via the disynaptic PC - cerebellar nuclei - IO feedback pathway (Marshall and Lang, 2009; Chaumont et al., 2013). Moreover, strong modulation of CS firing rates has been observed with direct cerebellar surface stimulation (Avlar et al unpublished data), suggesting that it is possible for ctACS to affect the state of the IO. One possible way to reconcile these results would be if the electric field distribution with ctACS is less concentrated than with direct surface stimulation, as is likely to be the case. As a result, PCs with strong SS modulation are less likely to be tightly clustered such that their activity would lead to strong modulation of the inhibitory cerebellar nuclear neurons that project to the specific IO region that projects back to the PC being recorded. Recordings of cerebellar nucleo-olivary neurons during ctACS are needed to address this issue more definitively. Another factor that might limit the modulation of CS firing rate by ctACS is the properties of the PC - cerebellar nucleo - olivary pathway (Best and Regehr, 2009; Najac and Raman, 2015), which should make it act like a low pass filter, and which would therefore limit the effective transmission of SS modulation to lower ctACS frequencies, which are relatively ineffective in driving SS activity (Kang et al., 2023; Mourra et al., 2025).

### Implications for neuromodulation strategies targeting cerebellar circuits

The distinct frequency dependencies of CS waveform and SS firing rate modulation could allow selective targeting of ctACS effects on plasticity. That is, while the effects of ctACS on CS waveform were similar across the range of tested frequencies (0.5 Hz - 50 Hz) (if anything slightly more prevalent at the lower end of that range), the strongest SS modulation is generated by 50-100 Hz ctACS. These distinct frequency dependencies may be particularly advantageous in rehabilitation contexts; a low ctACS frequency could be chosen to maximize plasticity within the cerebellar circuits, whereas a higher frequency could lead to more widespread, extracerebellar effects. Thus, the present findings emphasize the need for mechanistically informed and goal directed parameter selection when using ctACS.

## Author Contributions

EJL and MS developed the conceptual framework. DM and EJL designed the studies and drafted the initial manuscript. DM, EJL, and MS reviewed and revised the manuscript. DM collected and analyzed all SS and CS waveform datasets and produced data visualizations. EJL and DM created all data processing pipelines. JS performed the individual spikelet analyses. All the authors approved the final manuscript as submitted.

## Data Availability

The scientific data will be made freely available upon reasonable request, in compliance with NIH policies.

## References

Ahn, S., Mellin, J. M., Alagapan, S., Alexander, M. L., Gilmore, J. H., Jarskog, L. F., Fröhlich, F. (2019). Targeting reduced neural oscillations in patients with schizophrenia by transcranial alternating current stimulation. NeuroImage, 186, 126–136.

Asan, A. S., Sahin, M. (2019a) Modulation of Multiunit Spike Activity by Transcranial AC Stimulation (tACS) in the Rat Cerebellar Cortex. In: 2019 41st Annual International Conference of the IEEE Engineering in Medicine & Biology Society (EMBC), pp 5192–5195.

Asan, A. S., Gok, S., Sahin, M. (2019b). Electrical fields induced inside the rat brain with skin, skull, and dural placements of the current injection electrode. PLOS ONE, 14, e0203727.

Avlar, B., Rahman, R., Vendidandi, S., Cetinkaya, E., Asan, A. S., Sahin, M., Lang, E. J. (2023). Modulation of cerebellar cortical, cerebellar nuclear and vestibular nuclear activity using alternating electric currents. Front Syst Neurosci, 17, 1173738.

Bates, D., Mächler, M., Bolker, B., Walker, S. (2015). Fitting Linear Mixed-Effects Models Using lme4. Journal of Statistical Software, 67, 1–48.

Bazzigaluppi, P., De Gruijl, J. R., van der Giessen, R. S., Khosrovani, S., De Zeeuw, C. I., de Jeu, M. T. (2012). Olivary subthreshold oscillations and burst activity revisited. Front Neural Circuits, 6, 91.

Bell, C. C., Grimm, R. J. (1969). Discharge properties of Purkinje cells recorded on single and double microelectrodes. J Neurophysiol, 32, 1044–1055.

Best, A. R., Regehr, W. G. (2009). Inhibitory regulation of electrically coupled neurons in the inferior olive is mediated by asynchronous release of GABA. Neuron, 62, 555–565.

Bloedel, J. R., Roberts, W. J. (1971). Action of climbing fibers in cerebellar cortex of the cat. J Neurophysiol, 34, 17–31.

Burg, D., Rubia, F. J. (1972). Inhibition of cerebellar Purkinje cells by climbing fiber input. Pflugers Arch, 337, 367–372.

Burroughs, A., Wise, A. K., Xiao, J., Houghton, C., Tang, T., Suh, C. Y., Lang, E. J., Apps, R., Cerminara, N. L. (2017). The dynamic relationship between cerebellar Purkinje cell simple spikes and the spikelet number of complex spikes. J Physiol (Lond), 595, 283–299.

Chan, C. Y., Nicholson, C. (1986). Modulation by applied electric fields of Purkinje and stellate cell activity in the isolated turtle cerebellum. J Physiol (Lond), 371, 89–114.

Chan, C. Y., Hounsgaard, J., Nicholson, C. (1988). Effects of electric fields on transmembrane potential and excitability of turtle cerebellar Purkinje cells in vitro. J Physiol (Lond), 402, 751–771.

Chaumont, J., Guyon, N., Valera, A. M., Dugué, G. P., Popa, D., Marcaggi, P., Gautheron, V., Reibel-Foisset, S., Dieudonné, S., Stephan, A., Barrot, M., Cassel, J.-C., Dupont, J.-L., Doussau, F., Poulain, B., Selimi, F., Léna, C., Isope, P. (2013). Clusters of cerebellar Purkinje cells control their afferent climbing fiber discharge. Proc Nat Acad Sci, USA, 110, 16223–16228.

De Gruijl, J. R., Bazzigaluppi, P., de Jeu, M. T., De Zeeuw, C. I. (2012). Climbing fiber burst size and olivary sub-threshold oscillations in a network setting. PLoS Comput Biol, 8, e1002814.

Ebner, T. J., Bloedel, J. R. (1981a). Temporal patterning in simple spike discharge of Purkinje cells and Its relationship to climbing fiber activity. J Neurophysiol, 45, 933–947.

Ebner, T. J., Bloedel, J. R. (1981b). Correlation between activity of Purkinje cells and its modification by natural peripheral stimuli. J Neurophysiol, 45, 948–961.

Eccles, J. C., Llinás, R., Sasaki, K. (1966). The action of antidromic impulses on the cerebellar Purkinje cells. J Physiol (Lond), 182, 316–345.

Feng, T., Zhang, L., Wu, Y., Tang, L., Chen, X., Li, Y., Shan, C. (2024). Exploring the Therapeutic Effects and Mechanisms of Transcranial Alternating Current Stimulation on Improving Walking Ability in Stroke Patients via Modulating Cerebellar Gamma Frequency Band—a Narrative Review. The Cerebellum, 23, 1593–1603.

Fertonani, A., Ferrari, C., Miniussi, C. (2015). What do you feel if I apply transcranial electric stimulation? Safety, sensations and secondary induced effects. Clinical Neurophysiology, 126, 2181–2188.

Gilbert, P. F. (1976). Simple spike frequency and the number of secondary spikes in the complex spike of the cerebellar Purkinje cell. Brain research, 114, 334–338.

Granit, R., Phillips, C. G. (1956). Excitatory and inhibitory processes acting upon individual Purkinje cells of the cerebellum in cats. J Physiol (Lond), 133, 520–547.

Grimaldi, G., Argyropoulos, G. P., Boehringer, A., Celnik, P., Edwards, M. J., Ferrucci, R., Galea, J. M., Groiss, S. J., Hiraoka, K., Kassavetis, P., Lesage, E., Manto, M., Miall, R. C., Priori, A., Sadnicka, A., Ugawa, Y., Ziemann, U. (2014). Non-invasive Cerebellar Stimulation—a Consensus Paper. The Cerebellum, 13, 121–138.

Grimaldi, G., Argyropoulos, G. P., Bastian, A., Cortes, M., Davis, N. J., Edwards, D. J., Ferrucci, R., Fregni, F., Galea, J. M., Hamada, M., Manto, M., Miall, R. C., Morales-Quezada, L., Pope, P. A., Priori, A., Rothwell, J., Tomlinson, S. P., Celnik, P. (2016). Cerebellar Transcranial Direct Current Stimulation (ctDCS): A Novel Approach to Understanding Cerebellar Function in Health and Disease. The Neuroscientist, 22, 83–97.

Hankin, R. K. S. (2015). Circular Statistics in R. Journal of Statistical Software, 66, 1–3.

Ito, M. (2001). Cerebellar Long-Term Depression: Characterization, Signal Transduction, and Functional Roles. Physiological Reviews, 81, 1143–1195.

Ito, M., Simpson, J. I. (1971). Discharges in Purkinje cell axons during climbing fiber activation. Brain Research, 31, 215–219.

Jahnsen, H. (1986). Extracellular activation and membrane conductances of neurones in the guinea-pig deep cerebellar nuclei in vitro. J Physiol (Lond), 372, 149–168.

Kaiser, M., Wang, Y., Ten Oever, S., Duecker, F., Sack, A. T., Van De Ven, V. (2025). Simultaneous tACS-fMRI reveals state- and frequency-specific modulation of hippocampal-cortical functional connectivity. Communications Psychology, 3, 19.

Kang, Q., Lang, E. J., Sahin, M. (2023). Transsynaptic entrainment of cerebellar nuclear cells by alternating currents in a frequency dependent manner. Frontiers in Neuroscience, 17, 1282322.

Khaliq, Z. M., Raman, I. M. (2005). Axonal Propagation of Simple and Complex Spikes in Cerebellar Purkinje Neurons. J Neurosci, 25, 454–463.

Kuznetsova, A., Brockhoff, P. B., Christensen, R. H. B. (2017). lmerTest Package: Tests in Linear Mixed Effects Models. Journal of Statistical Software, 82, 1–26.

Lang, E. J., Tang, T., Suh, C. Y., Xiao, J., Kotsurovskyy, Y., Blenkinsop, T. A., Marshall, S. P., Sugihara, I. (2014). Modulation of Purkinje cell complex spike waveform by synchrony levels in the olivocerebellar system. Frontiers in Systems Neuroscience, 8, 210.

Latham, A., Paul, D. H. (1971). Spontaneous activity of cerebellar Purkinje cells and their responses to impulses in climbing fibres. Journal of Physiology (London), 213, 135–156.

Mano, N. (1970). Changes of simple and complex spike activity of cerebellar Purkinje cells with sleep and waking. Science, 170, 1325–1327.

Mardia, K. V. (1975). Statistics of Directional Data. J R Stat Soc B, 37, 349–371.

Marshall, S. P., Lang, E. J. (2009). Local changes in the excitability of the cerebellar cortex produce spatially restricted changes in complex spike synchrony. J Neurosci, 29, 14352–14362.

Maruta, J., Hensbroek, R. A., Simpson, J. I. (2007). Intraburst and Interburst Signaling by Climbing Fibers. The Journal of Neuroscience, 27, 11263–11270.

Mathy, A., Ho, S. S., Davie, J. T., Duguid, I. C., Clark, B. A., Hausser, M. (2009). Encoding of oscillations by axonal bursts in inferior olive neurons. Neuron, 62, 388–399.

McDevitt, C. J., Ebner, T. J., Bloedel, J. R. (1982). The changes in Purkinje cell simple spike activity following spontaneous climbing fiber inputs. Brain research, 237, 484–491.

Miyaguchi, S., Otsuru, N., Kojima, S., Saito, K., Inukai, Y., Masaki, M., Onishi, H. (2018). Transcranial Alternating Current Stimulation With Gamma Oscillations Over the Primary Motor Cortex and Cerebellar Hemisphere Improved Visuomotor Performance. Frontiers in Behavioral Neuroscience, 12, 132.

Monsivais, P., Clark, B. A., Roth, A., Häusser, M. (2005). Determinants of Action Potential Propagation in Cerebellar Purkinje Cell Axons. J Neurosci, 25, 464–472.

Mourra, D., Cavalieri, A. M., Casey, M. M., Sahin, M., Lang, E. J. (2025). Cerebellar Transcranial AC Stimulation produces a frequency-dependent bimodal cerebellar output Pattern. The Cerebellum, 24, 21.

Murphy, J. T., Sabah, N. H. (1970). The inhibitory effect of climbing fiber activation on cerebellar purkinje cells. Brain research, 19, 486–490.

Najac, M., Raman, I. M. (2015). Integration of Purkinje cell inhibition by cerebellar nucleo-olivary neurons. J Neurosci, 35, 544–549.

Naro, A., Bramanti, A., Leo, A., Manuli, A., Sciarrone, F., Russo, M., Bramanti, P., Calabrò, R. S. (2017). Effects of cerebellar transcranial alternating current stimulation on motor cortex excitability and motor function. Brain Structure and Function, 222, 2891–2906.

Priori, A., Ciocca, M., Parazzini, M., Vergari, M., Ferrucci, R. (2014). Transcranial cerebellar direct current stimulation and transcutaneous spinal cord direct current stimulation as innovative tools for neuroscientists: Cerebellar and Spinal tDCS. J Physiol (Lond), 592, 3345–3369.

Rasmussen, A., Jirenhed, D. A., Zucca, R., Johansson, F., Svensson, P., Hesslow, G. (2013). Number of spikes in climbing fibers determines the direction of cerebellar learning. J Neurosci, 33, 13436–13440.

Rowan, M. J. M., Bonnan, A., Zhang, K., Amat, S. B., Kikuchi, C., Taniguchi, H., Augustine, G. J., Christie, J. M. (2018). Graded Control of Climbing-Fiber-Mediated Plasticity and Learning by Inhibition in the Cerebellum. Neuron, 99, 999–1015 e1016.

Sato, Y., Miura, A., Fushiki, H., Kawasaki, T. (1992). Short-term modulation of cerebellar Purkinje cell activity after spontaneous climbing fiber input. J Neurophysiol, 68, 2051–2062.

Titley, H. K., Kislin, M., Simmons, D. H., Wang, S. S.-H., Hansel, C. (2019). Complex spike clusters and false-positive rejection in a cerebellar supervised learning rule. J Physiol (Lond), 597, 4387–4406.

Titley, H. K., Watkins, G. V., Lin, C., Weiss, C., McCarthy, M., Disterhoft, J. F., Hansel, C. (2020). Intrinsic Excitability Increase in Cerebellar Purkinje Cells after Delay Eye-Blink Conditioning in Mice. J Neurosci, 40, 2038–2046.

Yang, Y., Lisberger, S. G. (2014). Purkinje-cell plasticity and cerebellar motor learning are graded by complex-spike duration. Nature, 510, 529–532.

